# Antagonistic H3K79me-H3K9ac crosstalk determines elongation at housekeeping genes to promote pluripotency

**DOI:** 10.1101/2022.09.26.509534

**Authors:** Coral K. Wille, Xiaoya Zhang, Spencer A. Haws, John M. Denu, Rupa Sridharan

## Abstract

Pluripotent embryonic stem cells (ESCs) have a transcriptionally permissive chromatin environment enriched for gene activation-associated histone modifications as compared to somatic cells. A striking exception is DOT1L-mediated H3K79 methylation that is considered a positive regulator of transcription. Here we find that ESCs maintain low H3K79 methylation to facilitate RNA polymerase II (RNAPII) elongation for greater nascent transcription. Inhibiting DOT1L during the reprogramming of somatic to induced pluripotent stem cells (iPSCs) enables ESC-like RNAPII and transcriptional status. Mechanistically, DOT1L inhibition causes a local gain of histone acetylation at genes that lose the most H3K79me, which unexpectedly are ubiquitously expressed genes that perform essential functions in every cell, rather than lineage specifying genes. Maintenance of this elevated histone acetylation is required for the enhanced conversion to iPSCs upon DOT1L inhibition. Remarkably, increasing global DOT1L or site-specific tethering of DOT1L is sufficient to decrease H3K9ac in ESCs. We discover a high H3ac-low H3K79me epigenetic mechanism that promotes transcription elongation at ubiquitously expressed genes to enforce pluripotent cell identity.

## MAIN

During development, master transcription factors are necessary for lineage commitment because they control cell type-specific gene expression^1,2^. The introduction of such master regulators can even change the identity of already specified terminally differentiated cells^3^. For example, the transcription factors OCT4 (POU5F1), SOX2, KLF4, and c-MYC, are sufficient to reprogram somatic cells to induced pluripotent stem cells (iPSCs)^4^. Both lineage commitment and cell identity conversion are facilitated by a conducive epigenetic landscape^5–7^. Histone post-translational modifications (PTMs) are a primary determinant of the epigenetic environment and act through several broad mechanisms. These include directly affecting chromatin compaction by changing nucleosome surface charge and influencing the recruitment of chromatin remodeling enzymes to permit master regulator binding for RNA polymerase II (RNAPII) mediated transcription.

In contrast to differentiated cells, which require an accessible chromatin structure at lineage specific genes, pluripotent stem cells maintain their own identity while remaining poised for differentiation. These pluripotent properties are enabled by a less compacted chromatin structure^8^ whose features can be measured quantitatively by mass spectrometry for histone PTM abundance. Compared to unipotent somatic cells, pluripotent stem cells are depleted for histone PTMs associated with gene repression and enriched for those associated with gene activation, such as acetylation^9^. The open chromatin structure may allow embryonic stem cells (ESCs) to have a greater mRNA output per cell than somatic cells^10,11^, a phenomenon called hypertranscription^12^.

Paradoxical to this notion of a transcription ready chromatin structure, ESCs are severely depleted for histone H3K79 methylation (me)^9^. H3K79me is assumed to be a positive regulator of transcription because it is enriched on the gene bodies of highly and rapidly transcribed genes in somatic or transformed cell lines^13–15^. While histone modifications at promoters and enhancers reinforce cell specific transcriptional programs; whether gene body modifications, such as H3K79me2, are causal to cell identity is unexplored. DOT1L (disrupter of telomeric silencing 1-like) is the only enzyme that deposits H3K79 mono-, di-, and tri-methylation^16^. H3K79 methylation is thought to be removed by nucleosome turnover^17,18^. Importantly, despite their hypertranscriptional state, ESCs actively translocate DOT1L to the cytoplasm to maintain low nuclear levels^19^.

Knockout (KO) *Dot1l* ESCs continue to self-renew^20^ but are prone to totipotency^21^; and demonstrate compromised *in vitro* differentiation^22,23^. KO-*Dot1l* mice die at embryonic day 10.5_20_. Catalytic inhibition or depletion of DOT1L greatly increases cellular reprogramming efficiency from somatic to iPSCs^24,25^. Taken together, these findings indicate that low levels of DOT1L are conducive to, and even enhance pluripotency, but it is necessary for establishing somatic identity^26^.

Interestingly, and seemingly contradictory to the proposed role of H3K79 methylation as a positive regulator of transcription, compromising DOT1L leads to few steady-state transcriptional changes^23,25^. Thus the functional role of H3K79 methylation, as well as how it profoundly impacts cell fate change remain unknown. Here we show that ESCs maintain lower H3K79 methylation to increase the transition of RNA Polymerase II (RNAPII) into the gene body to promote nascent transcription. DOT1L inhibition during the reprogramming of somatic cells to iPSCs enhances nascent transcription and can even counteract the negative effects of paused transcription. By performing quantitative histone PTM mass spectrometry, we find a cell type specific response to DOT1L inhibition, with somatic cells increasing H3K27me3, ESCs increasing histone acetylation, and both modifications increasing in reprogramming populations. However, functionally, only maintaining the histone acetylation enhances reprogramming efficiency. Remarkably this increased histone acetylation primarily occurs at ubiquitously expressed genes that perform housekeeping functions rather than lineage-specific genes during the transition to iPSCs. The increased histone acetylation is located immediately downstream of the RNAPII pause site. We find that the balance between H3K79 methylation and H3K9 acetylation controls transcriptional elongation, a new kind of bivalent domain that can influence gene expression. DOT1L thus emerges as a master regulator of hyperacetylation and hypertranscription, two key properties of pluripotency, and implicates ubiquitously transcribed genes in cell fate decisions.

## RESULTS

### ESCs have lower H3K79me2 enrichment than somatic cells at shared genic locations

We were intrigued by the preference of pluripotent stem cells for low H3K79 methylation levels^20–23^. H3K79me2 was the most differential histone PTM between ESCs and mouse embryonic fibroblasts (MEFs) measured by quantitative mass spectrometry^9^. H3K79me2 levels in ESCs are also 3-5 fold lower in other somatic cells such as keratinocytes^25^ and astrocytes (Extended Data Fig. 1a). To gain insight into the genomic distribution of H3K79me2, we performed quantitative ChIP-seq with a spike in control^27^. In both MEFs and ESCs, 95% of H3K79me2 peaks are enriched on gene bodies (Extended Data Fig. 1b). As expected, MEF-specific and ESC-specific genes had a greater enrichment of H3K79me2 in the respective cell types (Fig. 1a-e). Surprisingly, 80% of H3K79me2 enriched genes overlap in both cell types (Fig. 1a) suggesting that the increased levels of this modification in MEFs do not occur at unique locations. Instead, ESCs have a lower enrichment of H3K79me2 at shared genes (Fig. 1b-c).

**Figure 1.**
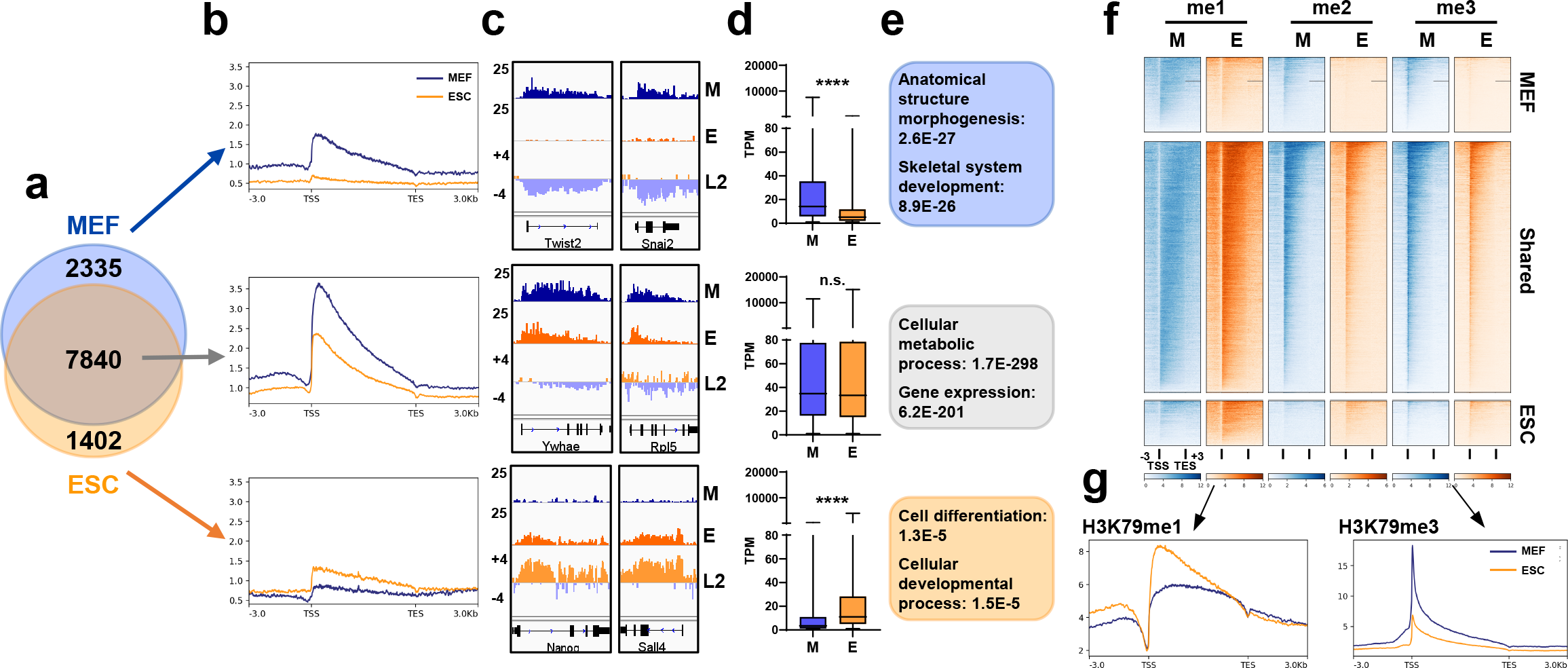
a. Overlap of genes with H3K79me2 peaks in MEFs (blue) and ESCs (orange). b-f: Genes with H3K79me2 peaks in MEF (M) unique, ESC (E) unique, and shared were assessed for: b. Relative H3K79me2 levels as metaplots. c. Example H3K79me2 IGV tracks (L2 = log2 fold change ESC/MEF). d. Expression (TPM). e. Gene ontology (GO). f. H3K79me1, H3K79me2, and H3K79me3 enrichment. g. Metaplot of H3K79me1 and H3K79me3 at shared genes with H3K79me2 peaks.

DOT1L is a distributive enzyme and the degree of H3K79 methylation depends on its local abundance^28^. We wondered whether H3K79 mono-or trimethylation, whose functions are not well-studied, might be elevated in the place of H3K79me2 at shared genes in ESCs. Like H3K79me2, ChIP-seq for H3K79me3 revealed a sharp peak at the transcription start site (TSS) of genes (Fig. 1f, Extended Data Fig. 1b-d) in both ESCs and MEFs with ∼3-fold greater enrichment in MEFs (Fig. 1g). In contrast to H3K79me2/me3, H3K79me1 showed a TSS enrichment in ESCs, and was evenly distributed across the gene body in MEFs (Fig. 1f-g). Altogether, our results indicate that the different degrees of H3K79 methylation have a cell type specific distribution pattern in pluripotent and somatic cells.

### ESCs are enriched for H3K79me1 instead of H3K79me2/me3 at highly expressed essential genes

We noticed that genes with shared H3K79me2 enrichment in both cell types were much more highly expressed than either MEF or ESC lineage specific genes (Fig. 1d). To further investigate how gene expression was influenced by the level of H3K79 methylation modification in each cell type, we binned genes into four groups based on steady-state RNA-seq measurements: High (>40 TPM), Middle (Mid) (12-40 TPM), Low (2-12 TPM), or Off (<2 TPM). Genes with shared H3K79me2 enrichment between ESCs and MEFs were expressed in both cell types (Extended Data Fig. 2a) and the majority (∼70%) remained within the same expression bin (Fig. 2a). In both MEFs and ESCs, higher expression correlated with total H3K79 methylation (me1+me2+me3) (Extended Data Fig. 2b). However, ESCs had significantly more H3K79me1 and less H3K79me2/3 at highly expressed genes as compared to MEFs (Fig. 2b-c, Extended Data Fig. 2c-e). The genes within the high bin in both MEFs and ESCs were enriched for functions in metabolic processes, gene expression, cell cycle, and intracellular transport that are considered essential for housekeeping in every cell type (Fig. 2a). Given this gene ontology, we examined gene expression data in several terminally differentiated cell types from distinct lineages such as keratinocytes, immune related B cells, and brain cells from the telencephalon and found that the shared H3K79me2 genes were always highly expressed (Fig. 2d).

**Figure 2.**
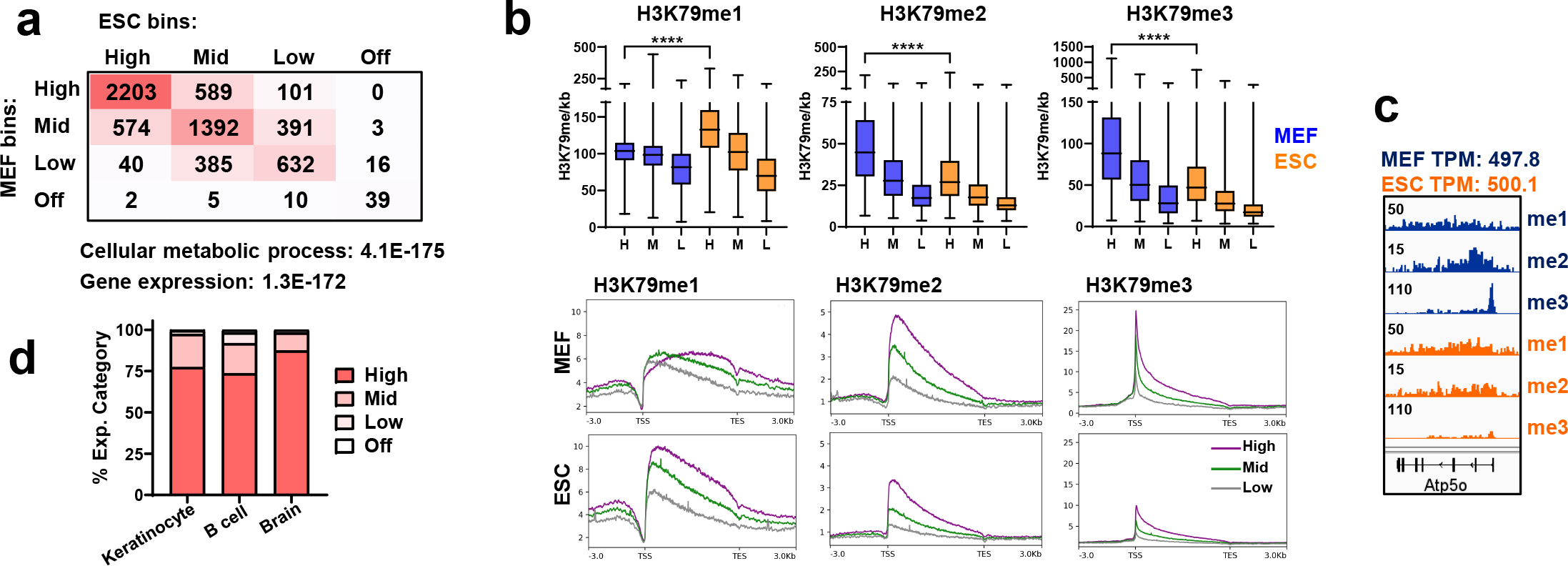
a. Comparison of expression category of genes with shared H3K79me2 peaks in ESCs vs. MEFs. GO of genes with high expression in both MEFs and ESCs displayed below the panel. b. Top: Normalized H3K79me genebody reads per kb gene length at High (H), Mid (M), and Low (L) expressed genes with shared peaks. ****P<0.0001 by unpaired two-tailed t-test. Bottom: Metaplots of H3K79me1/2/3 at High (Purple), Middle (Green), and Low (Gray) expressed genes with shared H3K79me2 peaks. c. H3K79me1/2/3 IGV tracks at a gene with shared H3K79me2 peaks and similar steady-state expression. d. Percent of MEF/ESC high (Fig. 2a, n = 2203) per expression category in other cell types: keratinocytes (Nefzger et al., 2017), immature splenic B cell (Aslam et al., 2021), and brain telencephalon (Franz et al., 2019).

Taken together, our analysis reveals that for ubiquitously expressed genes that have high expression in a multitude of tissues and perform housekeeping functions, enrichment of H3K79me1 is a better indicator of gene expression in ESCs as compared to H3K79me2/3 in MEFs. Furthermore, ESCs can maintain high steady state expression without H3K79me2/me3.

### ESCs have greater RNAPII elongation as compared to somatic cells

ESCs actively translocate DOT1L to the cytoplasm^19^ which likely prevents higher order H3K79me. Therefore, we investigated if low DOT1L and H3K79me2/3 is a feature or a facilitator of pluripotency. Steady state gene expression levels are dependent on the rates of transcription, mRNA processing, and degradation^29^. Transcription rate itself is determined by RNAPII initiation, release of RNAPII from pausing at about 50bp downstream of the TSS, and elongation through the gene body^30–34^. Since H3K79 methylation occurs on genes after transcription is initiated^35^, we hypothesized that cell type specific levels of H3K79 methylation influence RNAPII dynamics. Therefore, we performed quantitative ChIP-Seq for RNAPII in ESCs and MEFs, and interrogated the relative enrichment at the TSS compared to the gene body to determine the RNAPII traveling ratio (TR)^36^. ESCs display a significantly lower TR compared to MEFs at genes with shared H3K79me2 enrichment (Fig. 3a, list from Fig. 1a) even at genes that were in the high expression bin and ubiquitously expressed in many cell types (keratinocytes, brain, and B cells) (Extended Data Fig. 3a). ESCs have much greater protein levels of RNAPII (Fig. 4d) and the general transcription factor TFIID^37^ that promotes RNAPII initiation. Yet, unexpectedly, ESCs have a lower TR than MEFs because of less RNAPII enrichment at the TSS than the gene body (Fig. 3a).

**Figure 3.**
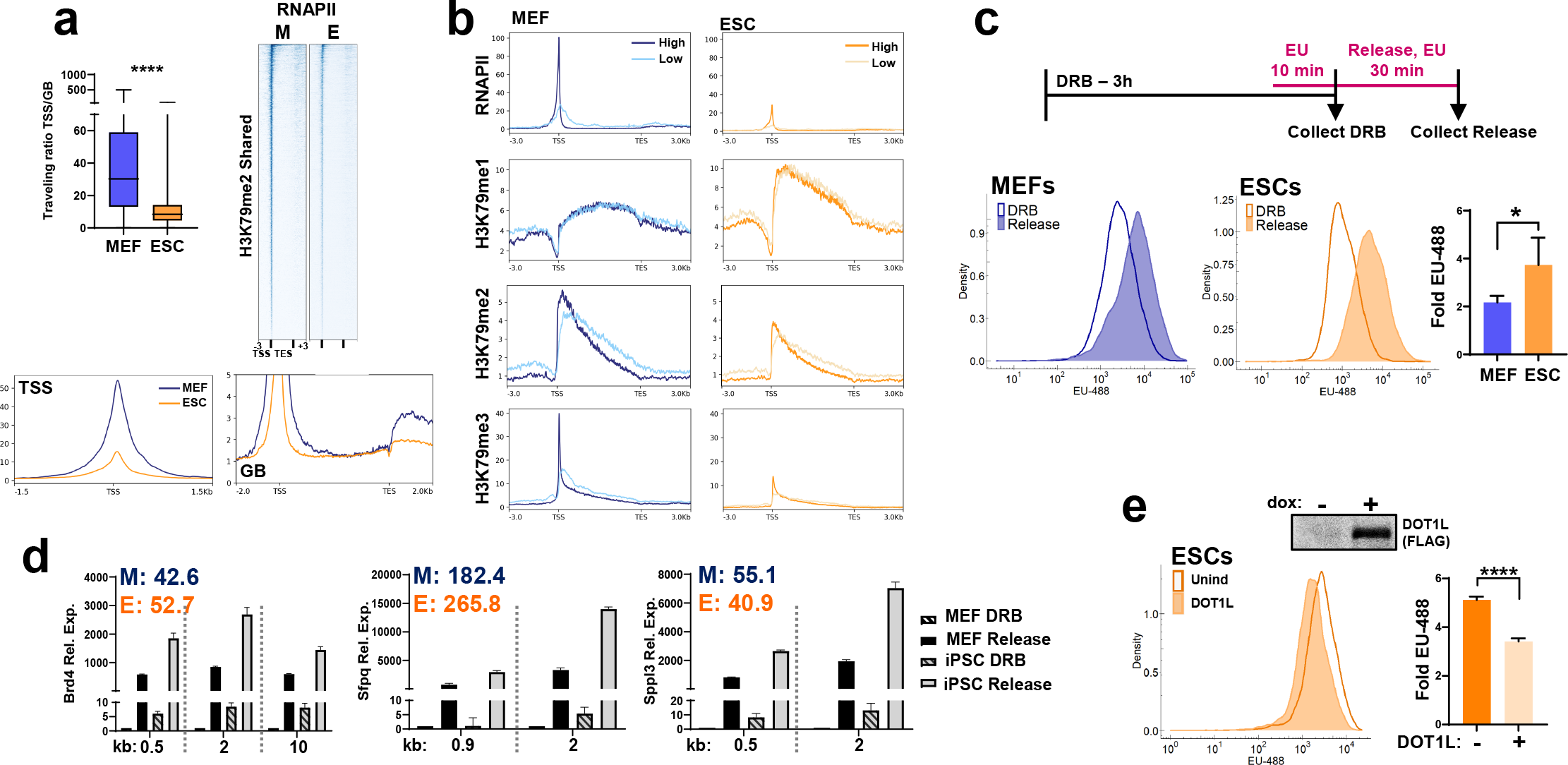
a. RNAPII analysis at shared genes with H3K79me2 peak (Fig. 1a) in MEFs (M) and ESCs (E) of - Left: Traveling ratio. Data are the mean (n = 2). ****P<0.0001 by unpaired two-tailed t test. Right: RNAPII heatmap. Bottom: RNAPII metaplots centered at the TSS or genebody (GB). b.Genes with a shared H3K79me2 peak, with high and similar expression (within 2 fold-TPM in MEFs and ESCs) were sorted by RNAPII traveling ratio. Metaplots of RNAPII, H3K79me1, H3K79me2, and H3K79me3 of the highest (dark) and lowest (light) travel ratio quartiles (n = 415 genes). c. Top: Nascent RNA experimental design. DRB was added for 3 hours to stop RNAPII transcription. 5-EU was added for the last 10 min of DRB treatment so that 5-EU nucleotide would be processed by the salvage pathway. The control DRB unreleased sampled was collected. Alternatively, DRB was removed by PBS washes to release RNAPII, and media containing fresh 5-EU was replaced. Transcription was allowed to proceed for 30 min before sample collection. Bottom: Flow cytometry EU-488 signal density plot of MEFs and ESCs. Outline = unreleased cells paused with DRB for 3 hours, Shaded = cells paused for 3 hours with DRB and then released for 30 min in the presence of 5-EU. Right: Fold increase of EU-488 signal after release, DRB sample set to 1. Data are the mean + S.D. (n = 3). *P<0.05 by ratio paired two-tailed t-test. d. Relative nascent RNA expression labeled with 5-EU for 30 min after DRB release, measured by RT-qPCR. Primers located in introns at the specified distance (kb) from the TSS. Unreleased DRB MEFs set to 1. Inset: Steady state expression (TPM), M = MEFs, E = ESCs. e. Top: Full-length FLAG-DOT1L dox-induced expression in ESCs. Left: Nascent RNA flow cytometry EU-488 signal density plot of ESCs released for 30 min, with and without DOT1L overexpression. DRB/5-EU treatment design as in Fig. 3c. Outline = Uninduced (Unind) control ESCs, Shaded = ESCs induced to express full length DOT1L for 3 days. Right: Fold increase of EU-488 signal after 30 min release, DRB sample set to 1. Data are the mean + S.D. (n = 3). ****P<0.0001 by unpaired two-tailed t-test.

**Figure 4.**
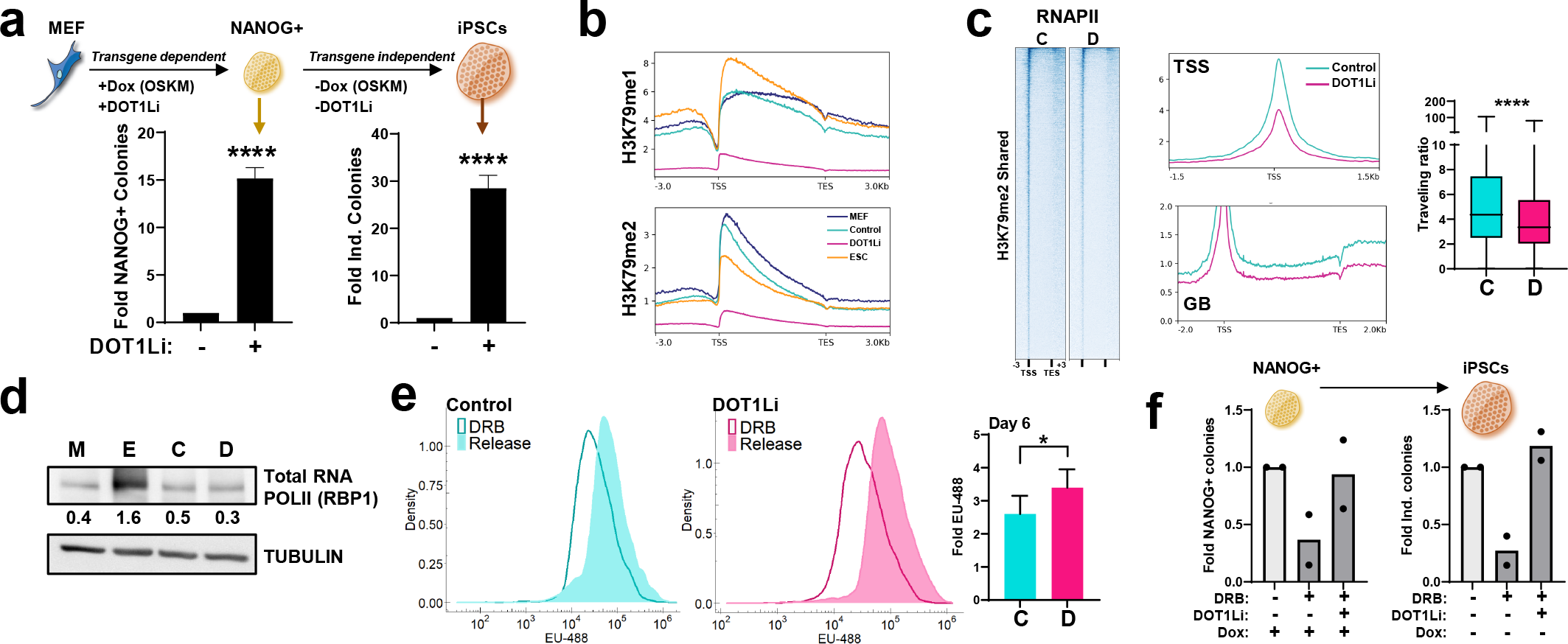
a.Top: Reprogramming scheme. OCT4, SOX2, KLF4, c-MYC (OSKM) is induced in MEFs with doxycycline (dox) in the presence of DOT1Li or DMSO control. Transgene dependent colonies are groups of at least 4 cells that express NANOG in dox on days 6-8. Transgene independent colonies maintain NANOG expression after dox and DOT1Li removal. Bottom: transgene dependent (Left) and independent (Right) NANOG+ colonies. Control treated cells set to 1. Data are the mean + S.D. (n = 3). ****P<0.0001 by unpaired two-tailed t test. b. Metaplot of H3K79me1 and H3K79me2 at genes with shared H3K79me2 peaks in MEFs and ESCs (from Fig. 1a). c. RNAPII analysis at shared genes with H3K79me2 peak in reprogramming day 4 control (C) and DOT1Li (D) treated cells of – Left: RNAPII heatmap. Middle: RNAPII metaplots centered at the TSS or genebody (GB). Right: Traveling ratio. Data are the mean (n = 2). ****P<0.0001 by unpaired two-tailed t test. d. Immunoblot of total RNAPII in MEFs (M), ESCs (E), and day 4 reprogramming of cells treated with Control (C), or DOT1Li (D). Relative quantitation shown below. e. Left: Flow cytometry EU-488 signal density plot of day 6 of reprogramming cells treated with control or DOT1Li. Outline = unreleased cells paused with DRB for 3 hours, Shaded = cells paused for 3 hours with DRB and then released for 30 min in the presence of 5-EU. Right: Fold increase of EU-488 signal after release, DRB sample set to 1. Data are the mean + S.D. (n = 3). *P<0.05 by ratio paired two-tailed t-test. f. Reprogramming of cells treated with RNAPII inhibitor DRB (dark gray), with and without DOT1Li, measured as transgene dependent (Left) and independent (Right) colonies. Cells treated with control set to 1. Biological replicates depicted as dots (n = 2).

We then asked how H3K79 methylation correlated with TR. In highly expressed genes with similar steady-state expression in both ESCs and MEFs, a low TR corresponded with low H3K79me2/me3 at the TSS (Fig. 3b). Interestingly, in ESCs, genes with a lower TR had a post-TSS shifted pattern of H3K79me1 and H3K79me2 (Fig. 3b), which could reflect a faster moving RNAPII that carries DOT1L along with it.

The decreased TSS-associated RNAPII as well as the gene body shifted H3K79 methylation signal suggests that in ESCs RNAPII transitions from pause to elongation more efficiently than MEFs. To directly measure the transition to elongation, RNAPII was paused with the addition of 5,6-Dichloro-1-beta-D-ribofuranosylbenzimidazole (DRB) then released in the presence of 5-ethynyl uridine (EU) for 30 minutes, to capture *de novo* synthesized RNA after pause release (Fig. 3c). By coupling EU incorporation to a fluorescent readout, we measured newly synthesized (nascent) RNA in single cells using flow cytometry. ESCs had significantly more nascent RNA (∼4-fold compared to DRB control) after DRB release as compared to MEFs (∼2 fold compared to DRB control) (Fig. 3c, Extended Data Fig. 3b).

To gain a gene-centric view of nascent RNA production, we chose three candidate loci that perform essential functions in widely different housekeeping pathways - chromatin remodeling (*Brd4*), splicing (*Sfpq*), and intracellular secretion (*Sppl3*) - which were highly and evenly expressed in MEFs and ESCs by steady state measurements (Fig. 3d). After pausing RNAPII with DRB, we measured EU-labeled nascent RNA by intronic RT-qPCR. At locations ranging from 0.5kb to 2kb or more from the TSS, we found that ESCs had a greater amount of nascent RNA at all three loci than MEFs (Fig. 3d, Extended Data Fig. 3c). Moreover, ESCs had more nascent RNA in the presence of DRB even before release indicating a reduced sensitivity to DRB and/or pause regulation (Fig. 3d, Extended Data Fig. 3c) in accordance with lower relative RNAPII occupancy at the TSS in ESCs (Fig. 3a).

To directly test the causal connection between *de novo* RNA synthesis and H3K79me2/me3, we globally increased the levels of DOT1L in ESCs. We integrated full-length DOT1L into a single locus in the genome that was controlled by a doxycycline promoter (Fig. 3e). After DRB-pause release there was a significant ∼2-fold reduction in newly synthesized RNA in the DOT1L-induced ESC when compared to control (Fig. 3e). Taken together, we demonstrate that ESCs hypertranscribe and contain more nascent RNA, because of enhanced RNAPII elongation that is controlled by DOT1L mediated H3K79 methylation.

### Erasure of H3K79 methylation promotes reprogramming by altering RNAPII dynamics

We next examined how DOT1L and H3K79me affect acquisition of the pluripotent state with its unique RNAPII profile. While DOT1L is dispensable for the maintenance of ESCs^20,22,23^, its catalytic activity is a barrier to the reprogramming of mouse and human somatic cells to iPSCs^24,25,38^. Our previous results have shown that DOT1L inhibition does not increase reprogramming efficiency by affecting lineage specific gene expression^25^. Therefore, we investigated whether reducing H3K79 methylation causally influenced this change in cell identity by altering RNAPII dynamics.

We induced reprogramming in MEFs that have a single transgenic copy of the *Oct4, Sox2, c-Myc* and *Klf4* (OSKM) integrated into the genome under a doxycycline promoter in conjunction with a small molecule pharmacological inhibitor of DOT1L catalytic activity SGC0946 (DOT1Li)^39^. This inhibitor lowers DOT1L association with chromatin and thus may function by reducing local enzyme concentration^40^. Upon DOT1Li, we observe a ∼15 fold increase in colonies expressing the pluripotency factor NANOG in the presence of OSKM transgene expression by day 6 (Fig. 4a). A stringent measure of *bona fide* iPSC formation is sustained pluripotency without exogenous OSKM transgene expression which we measure by withdrawing doxycycline and DOT1Li (Fig. 4a). Even though the total number of NANOG+ colonies decreases after the OSKM transgene is turned off (Extended Data Fig. 4a), DOT1Li increases transgene-independent stable iPSCs by 30-fold (Fig. 4a).

Using this system, we interrogated the localization of H3K79me1/2/3 upon DOT1Li. We performed ChIP-seq on day 4 of reprogramming, even before the appearance of transgene-dependent NANOG+ colonies (Extended Data Fig. 4b), to capture the molecular effects in the transition to pluripotency albeit in a heterogenous population. DOT1Li greatly reduced H3K79me1, me2, and me3 enrichment (Fig. 4b, Extended Data Fig. 4c). The remainder of the peaks after DOT1Li exposure were still genic and retained at genes with the highest expression (Extended Data Fig. 4d-e).

We then performed ChIP-Seq for RNAPII in reprogramming populations to determine transcriptional dynamics with reduced H3K79me. Remarkably, DOT1Li populations had lowered TR as compared to the control due to a decrease in RNAPII at the TSS (Fig. 4c), making the pattern of enrichment resemble that in ESCs (Fig. 3a). The gene body enrichment of RNAPII in DOT1Li exposed cells was also lower than in control reprogramming populations (Fig. 4c). This is likely because the total RNAPII protein does not reach the levels observed in ESCs by day 4 of reprogramming (Fig. 4d). Thus, removal of H3K79me during reprogramming allows cells to acquire a more pluripotency-like RNAPII enrichment pattern.

To uncover the functional consequence of this lowered TR, we measured nascent RNA production. DOT1Li-treated reprogramming populations had a ∼3.4-fold increase in total nascent transcripts as compared to ∼2.6-fold in control treatment (Fig. 4e, Extended Data Fig. 4f) after DRB-pause release. Even in a heterogenous reprogramming population with lower levels of RNAPII as compared to ESCs, DOT1Li leads to a greater accumulation of nascent RNA. We next tested whether this small increase in RNA synthesis facilitated pluripotency. While DRB is toxic at high concentrations, at lower concentrations it can be used to pause RNAPII without excessive cell death^41^. We performed reprogramming in the presence of non-toxic levels of DRB which yielded a reduced the number of iPSCs, indicating an increase in paused RNAPII is detrimental to pluripotency. Reprogramming efficiency was restored to control levels when DOT1Li was added in conjunction with DRB (Fig. 4f). Thus the negative effects of enforced RNAPII pausing on reprogramming can be counteracted by an increased transition to elongation via DOT1L inhibition. Taken together our results indicate that the reduction of H3K79 methylation promotes an ESC-like RNAPII transcription elongation and nascent RNA production which is required to enhance reprogramming to pluripotency.

### Low H3K79 methylation epigenetically enables higher H3K9ac

As we^25^ and others^22–24^ have found few DOT1L-mediated transcriptional effects on the regulation of single genes in pluripotency, we next interrogated if loss of H3K79me affects the epigenome to control nascent RNA production on a large-scale. While it is well established that transcriptional output is affected by histone modification, less is known about the epigenetic regulation of RNAPII processivity. We used mass spectrometry to quantify global levels of histone PTMs in an unbiased way in MEFs and ESCs subjected to DOT1Li. As expected in both cell types, there was an almost complete loss of H3K79me1/2 upon DOT1Li (Extended Data Fig. 5a). However, we found opposite effects on the repressive and activating histone PTMs in MEFs and ESCs exposed to DOT1Li (Fig. 5a). While the repressive H3K27me3 increased by statistically significant levels in MEFs, a collection of histone acetylations found at the promoters of active genes were significantly increased only in ESCs (Fig. 5a, Extended Data Fig. 5a-b).

**Figure 5.**
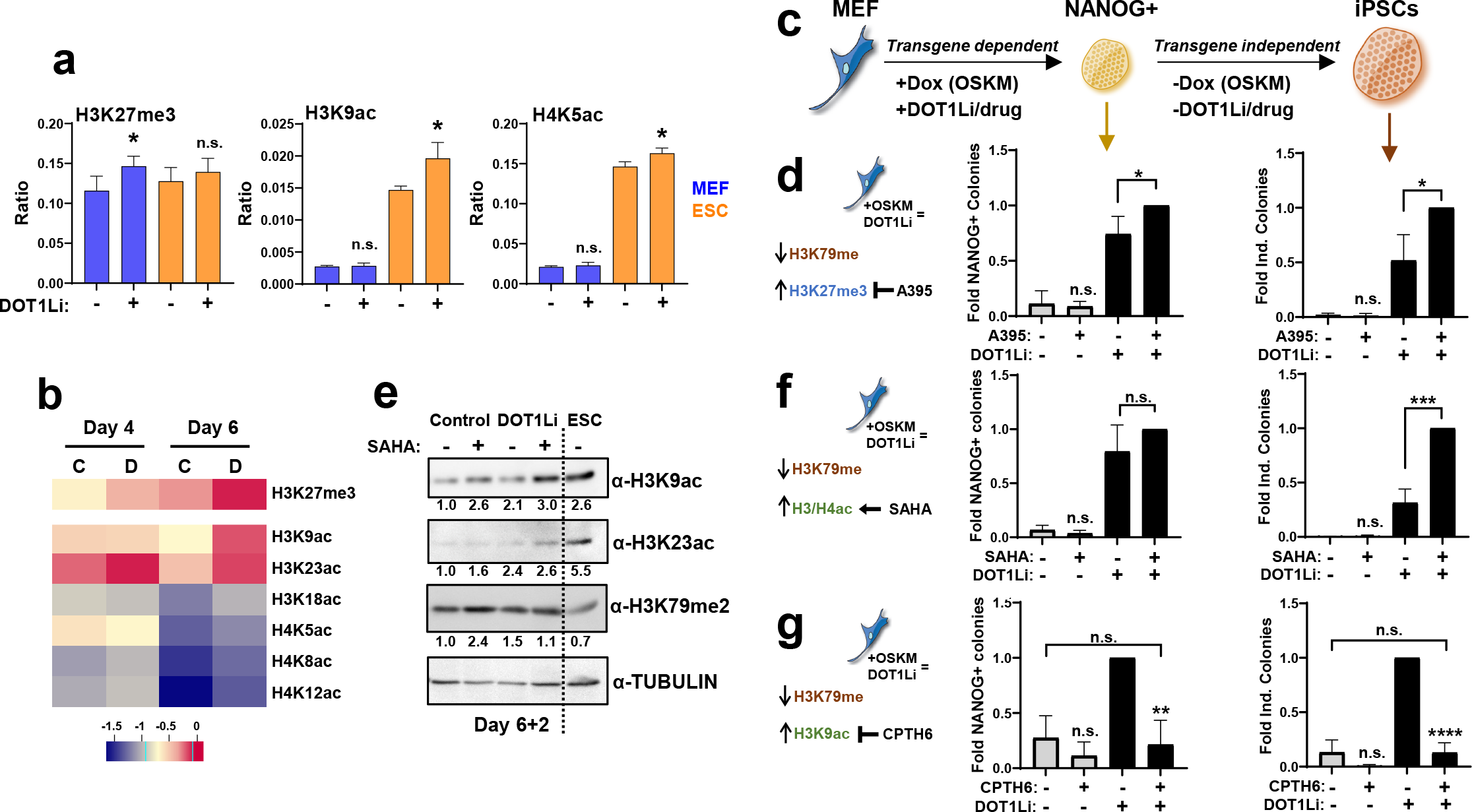
a. Average peptide ratio containing the specified modification determined by mass spectrometry of MEFs (blue) or ESCs (orange) treated with control (-) or DOT1Li (+) for 4 days. Significance determined relative to control treatment of the same cell type. Data are the mean + S.D. (n = 3-4). *P<0.05 or not significant (n.s.) P>0.05 by unpaired two-tailed t-test. b. Log2 fold change average peptide ratio heatmap of select modifications in control (C) and DOT1Li (D) treated reprogramming cells relative to the ratio modified in ESCs. Data are the mean (n = 3-4). c. Reprogramming scheme of transgene dependent and independent NANOG+ colonies. d. Left: Schematic of H3K27me3 targeting during reprogramming. Reprogramming of cells treated with control (gray) or DOT1Li (black), with and without EED inhibitor A395 measured as transgene dependent (Middle) and independent (Right) colonies. Cells treated with DOT1Li and A395 set to 1. Data are the mean + S.D. (n = 3). *P<0.05 or not significant (n.s.) P>0.05 by unpaired two-tailed t-test. e. Immunoblot 2 days post dox removal (after 6 days of reprogramming) of H3K9ac, H3K23ac, H3K79me2, and TUBULIN of control or DOT1Li cells treated with SAHA or vehicle beginning on day 3 of reprogramming. Relative quantitation shown below. f. Left: Schematic of histone H3 and H4 acetylation targeting during reprogramming. Reprogramming of cells treated with control (gray) or DOT1Li (black), with and without HDAC inhibitor SAHA measured as transgene dependent (Middle) and independent (Right) colonies. Cells treated with DOT1Li and SAHA set to 1. Data are the mean + S.D. (n = 3). ***P<0.001 or not significant (n.s.) P>0.05 by unpaired two-tailed t-test. g. Left: Schematic of H3K9ac targeting during reprogramming. Reprogramming of cells treated with control (gray) or DOT1Li (black), with and without GCN5 inhibitor CPTH6 measured as transgene dependent (Middle) and independent (Right) colonies. Cells treated with DOT1Li set to 1. Data are the mean + S.D. (n = 3). ****P<0.0001, **P<0.01, or not significant (n.s.) P>0.05 by unpaired two-tailed t-test.

Thus, H3K79me negatively regulates the deposition of repressive H3K27me3 or activating H3 and H4 acetylation marks, depending on the cell type.

In contrast to the cell-type specific effects, DOT1Li consistently increased both H3K27me3 and H3/H4 acetylation in reprogramming populations (Fig. 5b, Extended Data Figs. 5c-d, 6). We confirmed that multiple histone acetylation marks were indeed increased using siRNA as an orthogonal approach to deplete DOT1L during reprogramming (Extended Data Fig. 5e-g). Importantly, none of the histone modifying enzymes that have catalytic activity towards either H3K27 methylation or H3/H4 acetylation changed in RNA expression upon DOT1Li treatment (Extended Data Fig. 5h).

While previous work has shown that H3K79me2/3 blocks the spread of H3K27me3 in leukemia by chromatin immunoprecipitation^42^, a connection between H3K79 methylation and promoter histone acetylation is novel. Leukemia shares the infinite self-renewal phenotype of pluripotent stem cells. ESCs also have a hyperacetylated epigenome^9^ that enables their open chromatin structure. Therefore an increase in either H3K27me3 or in H3/H4 acetylations could potentially promote pluripotency. Hence we sought to determine which of the two different classes of epigenetic alterations that we observed above were functionally relevant to pluripotency acquisition (Fig. 5c).

H3K27me3 is deposited by the Polycomb Repressive Complex 2 which includes the EED component that mediates the spread of H3K27me3 to adjacent histones. We measured the effect of inhibiting EED in reprogramming in combination with DOT1Li (Extended Data Fig. 5i). Like our results above (Fig. 4a), DOT1Li increased reprogramming efficiency, while EEDi alone did not have any effect (Fig. 5d). However, when the DOTLi and EEDi were combined, we found that reprogramming efficiency was modestly increased in both OSKM-dependent NANOG+ colonies and fully reprogrammed transgene independent iPSCs (Fig. 5d). Countering the DOT1Li-mediated H3K27me3 increase with EED inhibition did not reverse the enhanced reprogramming efficiency (Fig. 5d). Thus, DOT1Li-mediated H3K27me3 increase is detrimental to pluripotency acquisition through reprogramming.

Several different histone acetyltransferases are responsible for the individual histone acetylations that resulted from DOT1Li. Therefore, to interrogate the broad increase in histone acetylation (Fig. 5b), we combined DOT1Li with the class I/II HDAC inhibitor SAHA to enhance retention of acetylation in the reprogramming population (Fig. 5e). The DOT1Li+SAHA combination did not cause any change in the number of OSKM-dependent NANOG+ colonies (Fig. 5f). Strikingly, upon withdrawal of doxycycline, there was a significant increase in the OSKM transgene independent stable iPSC in DOT1Li+SAHA treated reprogramming populations (Fig. 5f). DOT1Li induces a transient increase in H3/H4 acetylations, that are stabilized by using SAHA (Fig. 5e). Taken together, it is the retention of histone hyperacetylation triggered by the loss of H3K79me that specifically promotes transgene independence to generate *bona fide* iPSCs.

Among the acetylated histones, H3K9ac was most increased in both reprogramming populations (Fig. 5b) and in ESCs (Fig. 5a) upon DOT1Li. We next tested the effects of inhibiting the H3K9 acetyltransferase GCN5 (Extended Data Fig. 5j). Similar to depletion of GCN5^43^, its pharmacological inhibition with CPTH6 reduced both transgene dependent and independent iPSC colonies (Fig. 5g). This effect was much more dramatic when the GCN5 inhibitor was combined with DOT1Li suggesting that the gain of H3K9ac is especially relevant to the gain of pluripotency.

### Inhibition of DOT1L increases H3K9ac at ubiquitously expressed genes that have shared H3K79me enrichment in both ESCs and MEFs

Since histone acetylation increases upon H3K79me loss, and DOT1Li-mediated reprogramming is counteracted by GCN5 inhibition, we investigated where the gain of H3K9ac was localized. We performed ChIP-seq for H3K9ac on day 4 of reprogramming treated with control or DOT1Li and compared it to locations in MEFs and ESCs. Similar to H3K79 methylation, H3K9ac is overwhelmingly localized to promoter and genic locations (Extended Data Fig. 7a). Approximately 60% of genes with H3K9ac enrichment were shared in all four conditions/cell types (Extended Data Fig. 7b). Only 169 genes gained H3K9ac peaks exclusively in the DOT1Li reprogramming condition (Extended Data Fig. 7b) demonstrating that DOT1Li does not result in H3K9ac being installed at new locations during reprogramming.

Therefore, we quantitated the amount of H3K9ac in DOT1Li versus the control reprogramming population. We compared the features of the genes that gained H3K9ac (top 10% increased) and were relevant to reprogramming to the ones that lost H3K9ac (bottom 10% decreased) as a control (Fig 6a). The genes with the most increased H3K9ac upon DOT1Li shared H3K79me2 enrichment in both MEFs and ESCs rather than each cell type alone (Fig 6b). In this shared subset, the H3K9ac-increased (Inc) genes had a significantly higher level of H3K79me2 and paused RNAPII than the H3K9ac-decreased (Dec) genes in MEFs (Fig. 6c, Extended Data Fig. 7c), and were enriched for housekeeping functions (Fig. 6b). In contrast to the shared subset, there were no differences in the H3K79me2 levels between the H3K9ac increased and decreased genes in the exclusively MEF or ESC subset (Extended Data Fig. 7d). Taken together the direct effect of H3K79me2 loss in reprogramming is an increase of H3K9ac predominantly at genes with essential functions rather than lineage specifying genes.

**Figure 6.**
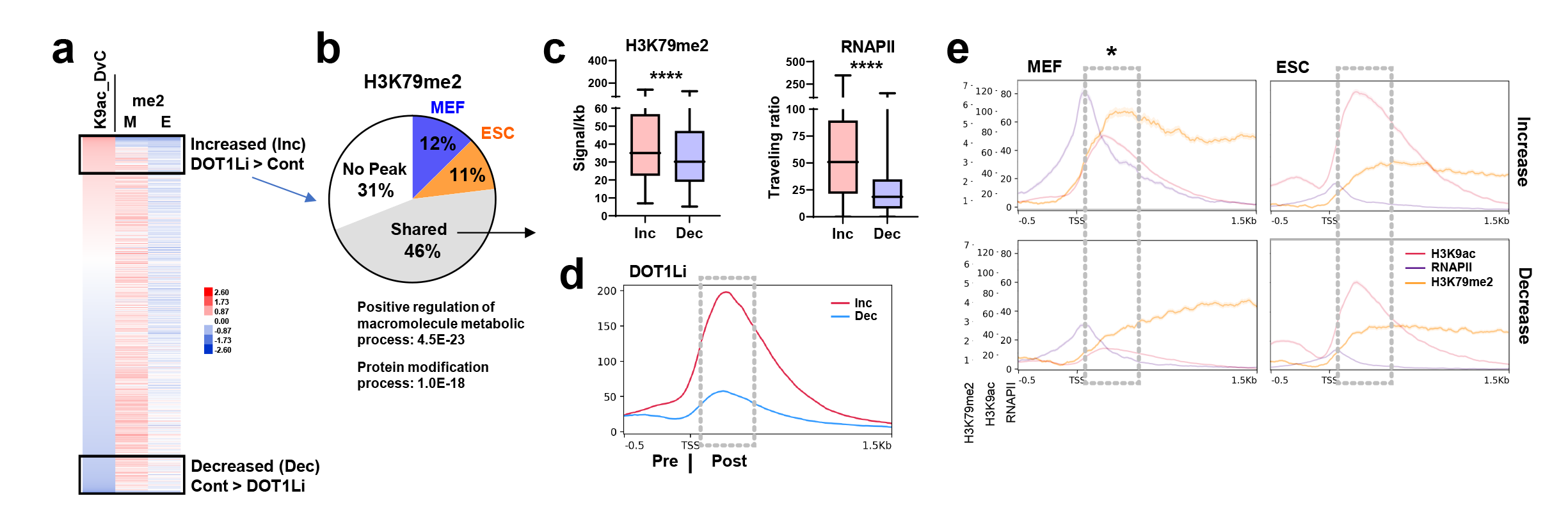
a. Heatmap of H3K79me2 (me2) enrichment of genes with an H3K9ac peak, sorted by relative H3K9ac enrichment (TSS to +1 kb) in day 4 reprogramming DOT1Li (D) vs. control (C). b. The H3K79me2 peak status of the top 10% of genes with increased H3K9ac in DOT1Li relative to control treated cells on day 4 of reprogramming. Bottom: gene ontology with a shared H3K79me2 peak in the top 10%. c-e. Genes with a shared H3K79me2 peak, in the top 10% with increased (Inc), compared to the bottom 10% with decreased (Dec), H3K9ac signal on day 4 of reprogramming in DOT1Li vs. Control were isolated for analysis of: c. Left: MEF H3K79me2 genebody signal per kb gene length. Right: MEF RNAPII traveling ratio (TSS/genebody). ****P<0.0001 by unpaired two-tailed t test. d. H3K9ac metaplot of day 4 reprogramming cells treated with DOT1Li. Area of increased H3K9ac boxed in gray. e. Overlaid metaplots of H3K79me2 (orange), H3K9ac (red), and RNAPII (purple) at H3K9ac increased (Top) and decreased (Bottom) genes. Area of increased H3K9ac boxed in gray. *Area of decreased H3K79me2. Note: Relative enrichment of modification is different and depicted by y-axes on left.

### Antagonism between H3K79me2 and H3K9ac proximal to the RNAPII pause site controls transcriptional elongation rate

In the DOT1Li reprogramming populations, we observed that the H3K9ac increased specifically after the TSS (Fig. 7d). To investigate this change in pattern of H3K9ac further, we compared MEFs and ESC relative to the RNAPII pause site. In MEFs at locations where H3K9ac is increased by DOT1Li, the apex of H3K79me2 enrichment coincided with that of H3K9ac after the RNAPII pause site, but not at the H3K9ac decreased locations. In contrast to the high H3K79me2/low H3K9ac in MEFs, ESCs had more elongating RNAPII coupled with a pattern of low H3K79me2/high H3K9ac. H3K9ac can act as a platform for recruiting the super elongation complex (SEC)^44^. Therefore, DOT1Li treatment during reprogramming switches the genes destined to have more elongating RNAPII, to an ESC-specific epigenetic profile. Together these results indicate that the relative levels of H3K9ac-H3K79me2 proximal to the RNA polymerase pause site could act as switch to promote transcriptional elongation.

**Figure 7.**
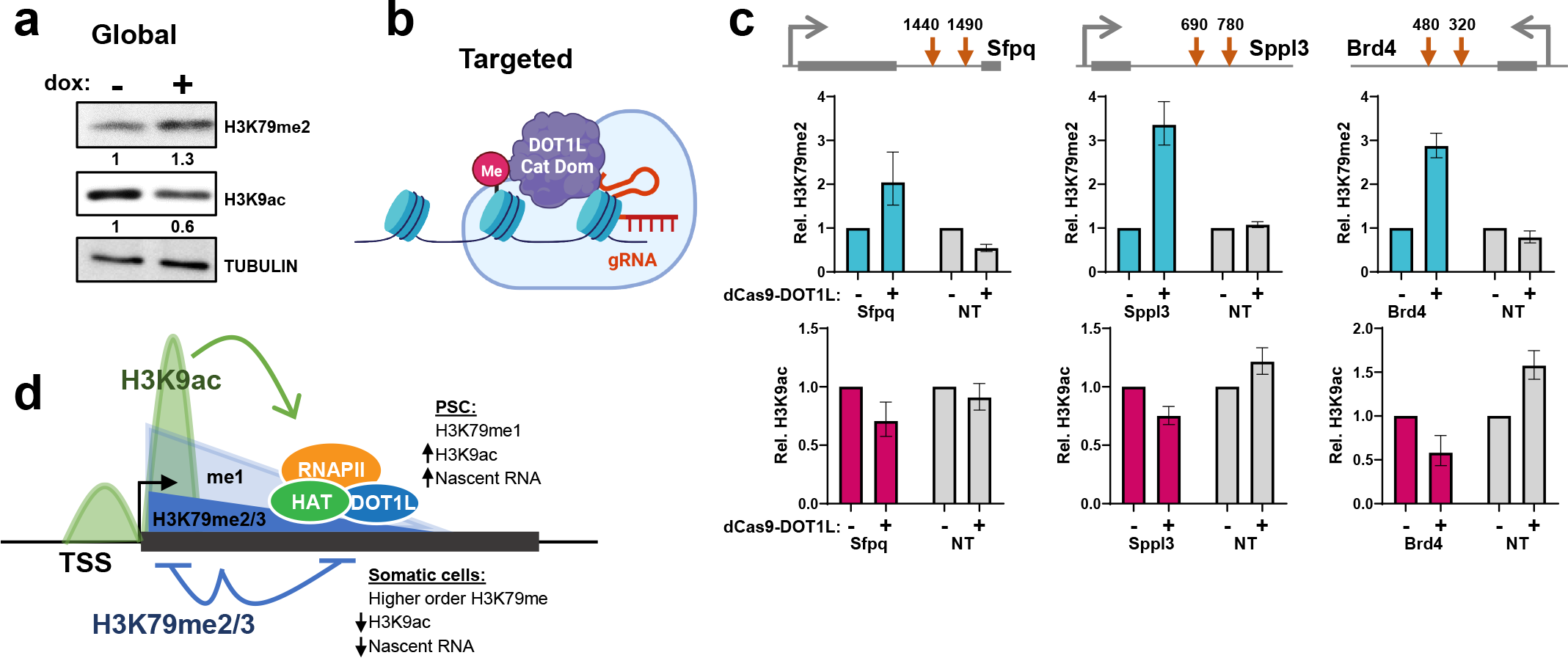
a. H3K79me2 and H3K9ac immunoblot of ESCs 48 hours after full-length DOT1L induction (+) compared to control uninduced (-) cells. Relative quantitation shown below. b.H3K79me targeting strategy: Dead-Cas9 (dCas9) fused to the DOT1L catalytic domain under the control of a doxycycline inducible promoter was integrated into the genome of ESCs. Cells were transduced with lentiviruses encoding small guide RNAs (gRNAs) and RFP to monitor efficiency of infection. ChIP-qPCR was performed 48h after dCas9-DOT1L induction. c. Top: The location of gRNAs at 3 genes targeted by dCas9-DOT1L to the apex of the H3K79me2 peak. H3K79me2 (Middle) and H3K9ac (Bottom) ChIP-qPCR at the targeted genes as well as non-targeted (NT) locations, relative to spike-in chromatin. ChIP was performed in cells infected with gRNA lentiviruses, with (+) and without (-) dCas9-DOT1L induction. d. Model of the role of H3K79 methylation in cellular identity and transcriptional regulation. DOT1L is recruited to transcribing genes. ESCs have higher levels of RNAPII and lower levels of DOT1L enabling a feed forward loop of faster elongation and enrichment of H3K9ac downstream of the TSS. Somatic cells are more biased to RNAPII initiation/pause detaining DOT1L at promoters resulting in higher order H3K79me2/3 because it is a distributive enzyme. Reduction of H3K79 methylation during reprogramming with DOT1Li allows establishment of ESC like H3K9 acetylation and greater *de novo* transcription.

To directly test this hypothesis, we induced full length DOT1L in ESCs. Strikingly, global H3K9 acetylation was reduced by 0.6-fold (Fig. 7a), even though H3K79me2 only increased 1.3-fold and did not reach the levels present in MEFs^9^. Notably none of the enzymes responsible for modulating H3 acetylation changed in expression at the RNA level (data not shown), strongly suggesting epigenetic means of regulation of these two modifications. Given that the antagonistic effects of H3K79me/H3K9a could still occur indirectly, we targeted the catalytic domain of DOT1L fused to nickase-deficient CAS9 to the H3K79me2 peak of specific loci (Fig. 7b). We chose loci that had similar steady state levels of transcription in MEFs and ESCs (Fig. 3d) that had housekeeping functions in different pathways, *Brd4, Sfpq*, and *Sppl3* (Fig. 7c). ChIP-qPCR confirmed an increase in H3K79me2 at the targeted locations (Fig. 7c, Extended Data Fig. 7e). As compared to control non-targeted locations, we observed a reduction in H3K9ac enrichment (Fig. 7c, Extended Data Fig. 7f). Thus a local increase in H3K79me2 can directly decrease H3K9ac.

## DISCUSSION

Taken together, DOT1L-mediated H3K79 methylation regulates an epigenetic switch of transcription associated histone modifications to promote a change in cell identity. H3K79me2/me3 in somatic cells acts as barrier to elongation (Fig. 7d), which is lowered in ESCs (Fig. 1b), leading to greater nascent RNA synthesis enabled by higher levels of RNAPII protein (Fig. 4d). The greater residence time of RNAPII at the TSS in somatic cells prolongs DOT1L association^45^, converting H3K79me1 to higher order methylations (Fig. 7d). This buildup of H3K79me2/3 at the TSS functions as a block of H3 acetylation gain that is essential to pluripotency. Our findings provide the first evidence for epigenetic control of the balance between transcription initiation and elongation, and clearly implicate that maintenance of a low H3K79me - high H3/H4ac domain is essential in reprogramming to pluripotency. Pluripotent cells have a higher demand for the products of housekeeping genes because of their greater biosynthetic requirements^12,46^. Our work emphasizes that histone modification dynamics within the gene bodies of non-lineage specific, essential genes plays an important role in the establishment and maintenance of pluripotency.

While *Dot1l* is expressed at similar transcript levels between somatic cells and ESCs^25^, DOT1L protein is shuttled to the cytoplasm by CDK1 mediated phosphorylation in ESCs^19^, thereby lowering nuclear H3K79me2/me3. Any further lowering of H3K79me - for example, by the deletion of *Dot1l* - has a minor effect on transcriptional elongation and is observed only in conjunction with a SEC inhibitor in ESCs^23^. In fact, H3K79me1 can explain half of the DOT1L phenotype in pluripotency acquisition^47^. Thus, there may be limits on how much further RNAPII dynamics can be altered by depleting H3K79me2/me3 to affect a phenotype.

Histone hyperacetylation is tightly connected to pluripotent identity as enforced maintenance of acetylation prevents differentiation^48^. Stabilizing the transiently induced promoter histone hyperacetylation by DOT1L inhibition is functionally important to achieving pluripotency (Fig. 5f). The distinct post-TSS pattern of H3K9ac gain could be due to decreased residence time of RNAPII and SAGA complex at the promoter region^49^ or DOT1L may counter recruitment of the SAGA complex as observed in erythroleukemia cells^50^. H3K9ac promotes pause release at candidate loci in HeLa cells by recruiting the SEC^44^. Our data are consistent with a feed forward loop where DOT1Li increases H3K9ac which synergizes with decreased H3K79me to enhance the RNAPII elongation (Fig. 7d). In addition to effects on RNAPII dynamics, histone hyperacetylation may facilitate reprogramming factor binding at the promoter^51^ to promote conversion to iPSCs.

DOT1L does not affect the ability of RNAPII to enter a pause in ESCs^23^. However, transcriptional output is determined more by exit rather than entry to a pause^11^. Models of transcriptional bursting have also shown that once a burst is initiated, pause release rather than RNAPII recruitment determines transcriptional output^52^. Thus pause release may be the important step that is regulated by H3K79me levels. Importantly, housekeeping genes in particular have been shown to be regulated by RNAPII pausing in ESCs^53^. Strikingly, the reduction of DOT1L can replace the reprogramming factor c-Myc^24^, which enhances RNAPII pause release^36^. This phenomenon seems to be evolutionarily conserved since in *C. elegans*, the deletion of *Zfp-1*, the catalytic activator of Dot-1, increased transcriptionally engaged RNAPII by GRO-seq and reduced the pausing index^54^.

Cellular environment may dictate cell fate after removal of H3K79 methylation such that gain of hyperacetylation and hypertranscription may be neither possible nor favorable in non-pluripotent or transformed cell types. In fact, we investigated the impact of DOT1Li on transdifferentiation between different types of somatic cells. We transdifferentiated MEFs into induced neurons (iNeuron) by the expression of *Brn2, Ascl1*, and *Myt1l* (BAM)^3^. iNeuron formation was significantly reduced by DOT1Li treatment (Extended Data Fig. 7g-i) indicating that pluripotency is uniquely compatible with low H3K79 methylation but differentiated cells are dependent on functional DOT1L. Furthermore, DOT1L deletion reduces transcription initiation in MEFs subjected to UV damage^55^, and in erythroleukemia cells^50^ without increasing nascent transcription. H3K79me may participate in an alternative epigenetic networks in somatic cells by regulating spread of H3K27me3 (Fig. 5a) and H3K27ac^42,56–60^.

Together our data provide insights into a new facet of pluripotency that focus on controlling the regulation of ubiquitously expressed genes in cell fate transitions, and open up new avenues for further improvement in reprogramming efficiency by targeting housekeeping genes in addition to lineage defining master regulators. Beyond cell fate transitions, our discovery of the antagonism between H3K79me and H3 acetylation has implications for cancers in which DOT1L protein levels are altered for combination therapy targeting both epigenetic modifications.

## METHODS

### Cell isolation and culture

Mice were maintained in agreement with our UW-Madison IACUC approved protocol. Male and female reprogrammable MEFs were isolated from embryos homozygous for the *Oct4*-2A-*Klf4*-2A-IRES-*Sox2*-2A-*c-Myc* (OKSM) transgene at the *Col1a1* locus and heterozygous for the reverse tetracycline transactivator (rtTA) allele at the *Rosa26* locus on day E13.5, as previously described^61^. MEFs were cultured in MEF media (DMEM, 10% FBS, 1x non-essential amino acids, 1x glutamax, 1x penicillin/streptomycin, and 4 μl/500 ml 2-Mercaptoethanol). Feeder MEFs were isolated at day E13.5 from DR4 embryos genetically resistant to geneticin (G418), puromycin, hygromycin, and 6-thioguanine. Feeder embryos were expanded for 3 passages and irradiated with 9000 rad. V6.5 ESCs and 2D4 iPSCs were grown on feeder MEFs in gelatinized dishes with ESC medium (knock-out DMEM, 15% FBS, 1x non-essential amino acids, 1x glutamax, 1x penicillin/streptomycin, 4 μl/525 ml 2-Mercaptoethanol, and leukemia inhibitory factor). ESCs and iPSCs were MEF-depleted by incubation for 30 min in a non-gelatinized dish before plating. Primary astrocytes were grown in DMEM with 1% FBS, 1x non-essential amino acids, 1x glutamax, and 1x penicillin/streptomycin, and isolated as previously described^38^. 293T and normal human dermal fibroblasts were cultured in DMEM and 10% FBS and acquired from ATCC.

### Inducible full length and targeted DOT1L ESC lines

DOT1L was cloned into pBS33 which can be integrated with FLP recombinase into the *Col1a1* locus of V6.5, under the control of a doxycycline (dox) inducible promoter^62^. For overexpression studies, the CDS encoding 5’-FLAG-tagged full-length mouse DOT1L protein was integrated. For DOT1L targeting studies, HA-tagged dead(d)-Cas9 in frame with the CDS of minimal human DOT1L (amino acids 2-416)^63^ was integrated. DOT1L expression was induced with 2 ug/ml of dox.

### Induced neuron transdifferentiation

MEFs that were homozygous for reverse tetracycline transactivator (rtTA) allele at the *Rosa26* locus were transduced with Tet-inducible *Brn2* (Addgene, 27151), *Ascl1* (Addgene, 27150), and *Myt1l* (Addgene, 27152) lentiviruses. MEFs were plated onto a coverslip at a density of 120,000 cells per 12 well, 2 days post transduction. Transdifferentiation was initiated with 2 μg/ml of doxycycline as previously described^3^, with control DMSO or 5 μM SGC0946 (ApexBio, A4167). Media was replaced day 3 post-induction with N3 media: DMEM/F12, 1x glutamax, 1x penicillin/streptomycin, 5 μg/ml insulin, 10 ng/ml FGF, and 1 x N-2 (ThermoFisher, 17502048) with and without SGC0946. Media was replaced every 2 days until coverslips were fixed for immunofluorescence on days 6-7.

### Lentiviral packaging and transduction

Lentiviral transfer vectors were transfected into 293T cells with packaging vector pspax2 (Addgene, 12260) and envelop vector vsvg using linear polyethylenimine. Media was changed to MEF media with 20 mM HEPES 4 hours post transfection. Virus containing media was harvested at 48h and 72h, combined, and filtered through a 0.45 μm PVDF filter. Virus containing media was combined with fresh media at a ratio of 1:1 and 10 μg/ml Hexadimethrine Bromide (polybrene) to transduce target cells.

### Reprogramming

Reprogrammable MEFs were plated at a density of 30,000 to 50,000 cells per gelatinized 12-well at day 0. Reprogramming was initiated with 2 ng/ml of doxycycline in ESC media containing DMSO control or drug treatment. SGC0946 (DOT1Li) was used at 5 μM (ApexBio, A4167), Vorinostat (SAHA) at 1 uM (Cell Signaling Technology, 12520S), CPTH6 hydrobromide at 40 uM (Cayman Chemical, 19828), 5,6-Dichloro-1-β-D-ribofuranosylbenzimidazole (DRB) at 10 uM (Sigma-Aldrich, D1916), and A-395 hydrochloride at 1 uM (Millipore Sigma, SML1923). Feeder MEFs were added at 50% confluency within days 0-2. Media containing fresh drugs and doxycycline was changed every 48 hours until efficiency was measured by immunofluorescence for the pluripotency factor NANOG. Days 6-8 were chosen as the endpoint as reprogramming proceeded long enough for stable colony formation, but not so long that cells became overcrowded in SGC0946/DOT1Li treatment. OSKM independent colonies were assessed by removing doxycycline and drugs for 2-4 days followed by immunofluorescence for sustained NANOG expression. A reprogrammed colony was considered as a grouping of at least 4 NANOG+ cells.

### Reprogramming statistical analysis

Reprogramming efficiency was calculated as the mean of three independent biological replicate experiments, each consisting of three technical replicate wells. Error bars depict the standard deviation (S.D.) of the three biological replicates unless otherwise noted. Individual replicate information is included in all legends. All reprogramming experiment significance was calculated using unpaired two-tailed t tests in Graphpad Prism 9 and P values are explained in legends.

### Immunofluorescence

Coverslips were fixed in 4% paraformaldehyde/PBS, permeabilized in 0.5% Triton-X/PBS, and washed in 0.2% Tween-20/PBS for 10 min each. Coverslips were blocked in Blocking Buffer (5% goat serum, 1x PBS, 0.2% Tween-20, 0.2% fish skin gelatin) for 30 min. Coverslips were stained with anti-NANOG (1:1000; Cell Signaling Technology, 8822S) or anti-TUJ1 (1:500; Novus Biologicals, MAB1195) in blocking buffer for 1 hour and then washed twice in wash buffer. Coverslips were stained with secondary goat anti-IgG-Dylight 488 (1:1000; ThermoScientific, 35552) in blocking buffer for 1 hour. Coverslips were washed 1x in wash buffer, 1x in wash buffer containing 0.1 μg/ml 4′,6-Diamidino-2-phenylindole dihydrochloride (Millipore Sigma, D8417), and 1x in wash buffer for 5 min each, before mounting on slides with aqua-polymount (Fisher Scientific, NC9439247). Counts and imaging were performed on Nikon Eclipse Ti using NIS Elements software.

### siRNA transfection

Cells were transfected every 48 hours in 0.5 mL per 12-well. 1 ul of DharmaFECT 1 (Fisher Scientific, T200104) was diluted in 49 ul of serum free DMEM and incubated for 5 min. The transfection reagent was then added to siRNA in 50 ul of serum-free DMEM and allowed to incubate for 20 min. Cells were transfected with 20 nmol of siRNA on days 0 and 2 of reprogramming, and then 40 nmol on day 4 to account for increased cell number. siRNA was purchased from Dharmacon (horizon): si*Dot1l* (J-057964-12) and non-targeting control (D-001810-01).

### RT-qPCR

RNA was isolated from cells on with the Isolate II RNA Mini Kit (Bioline, BIO-52702). 1 μg was converted to cDNA with qScript (Quanta, 95047) and 20 ng of cDNA (based on the original RNA concentration) were used for qPCR analysis in 10 ul reactions with SYBR Green (Bio-Rad, 1725124). Expression was calculated relative to the geometric mean of the two housekeeping genes.

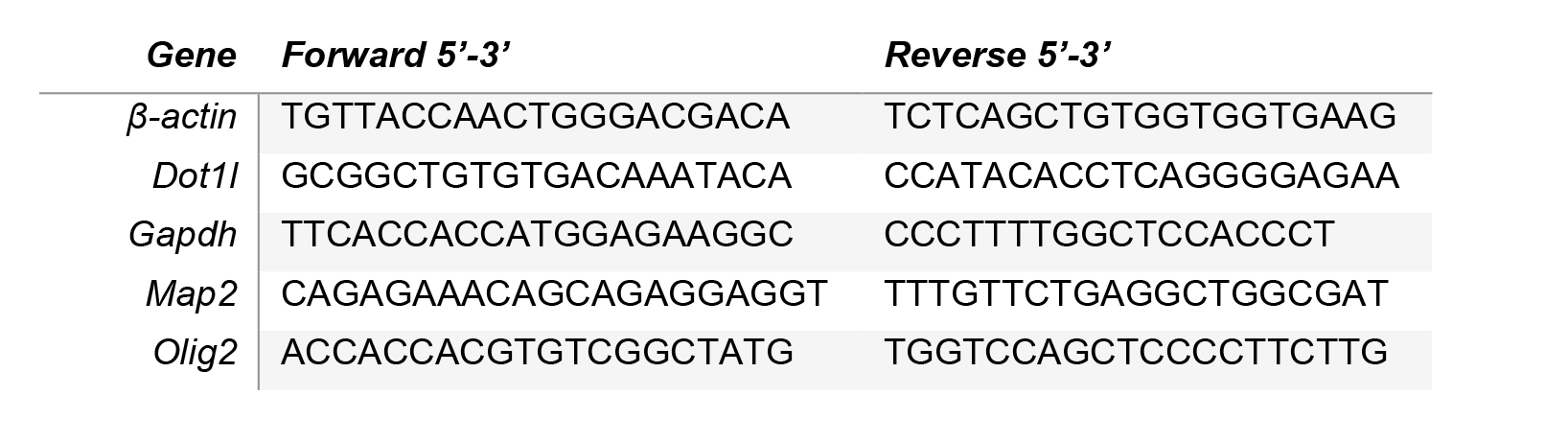

### Immunoblot

Cells were lysed in SUMO buffer: ¼ part I (5% SDS, 0.15 M Tris-HCl pH 6.8, 30% glycerol), ¾ part II (25 mM Tris-HCl pH8.3, 50 mM NaCl, 0.5% NP-40, 0.5% deoxycholate, 0.1% SDS), and 1x cOmplete protease inhibitors (Roche, 4693132001). Lysates were sonicated with a microtip for 5 sec at 20% amplitude and quantitated in DC Protein Assay kit II (BioRad, 5000112) against a BSA standard curve. 25 ug were loaded for H3K79me1/2 blots, 30 ug for RNAPII blots, 10 ug for acetylation blots, and 10 ug for H3K27me3. Protein was transferred to a nitrocellulose membrane, and blocked in 5% milk/PBS 1% Tween-20 for 30 min. Membranes were probed 1 hr to overnight in primary antibody, including: H3K79me1 (1:1000; Abcam, ab2886), H3K79me2 (1:1000; Active Motif, 39143), H3K9ac (1:1000; Active Motif, 39917), H3K23ac (1:1000; Active Motif, 39131), H3K27me3 (1:1000; Cell Signaling, 9733S), RNAPII RPB1 NTD D8L4Y (1:1000; Cell Signaling, 14958S), and α-TUBULIN (1:3000; Cell Signaling, 3873). Membranes were washed twice in Wash Buffer (PBS, 1% Tween-20) and incubated with secondary antibody for 1 hour. Membranes were washed 3x in Wash Buffer and imaged with ECL on ImageQuant LAS 4000. Tiff files were quantitated with Image Studio Lite V5.2 using the Add Rectangle function.

### Histone acid extraction and mass spectrometry

Two independent biological replicate expansions, each consisting of 4 technical replicates were analyzed. 4 million cells were resuspended in 800 ul Buffer A (10 mM Tris pH 7.4, 10 mM NaCl, 3 mM MgCl_2_) containing fresh 10 mM nicotinamide, 1 mM sodium butyrate, 4 uM trichostatin A, and 1x protease inhibitor (Roche, 4693132001), and transferred to a dounce homogenizer. Nuclei were isolated with 60 strokes using a tight pestle. The dounce was washed with 200 ul of buffer A and combined to make a total of 1 ml of lysate which was centrifuged at 800xg for 10 min at 4 C. The nuclei pellet was washed twice in cold PBS and spun as above. The pellet was resuspended in 500 ul of 0.4 N H_2_SO_4_, and rotated for 4 hours at 4C. Tubes were then spun at 3400xg for 10 min at 4C, and the supernatant containing histones was isolated. 125 ul of 100% TCA was added and incubated overnight. Histones were precipitated by centrifugation at 3400xg for 5 min at 4C and washed twice with 100% ice cold acetone. The pellet was air dried for 4 hours and then resuspended in 100 ul H2O. The solution was centrifuged at 3400xg for 2 min and the supernatant was collected. Protein concentration was measured against a standard curve with DC Protein Assay kit II (BioRad, 5000112) assay. 5 ug of protein were dried by speedvac and prepared for mass spectrometry as previously described^64^. Peptides were analyzed with Epiprofile 2.0^65^.

### Chip-Seq and library

At least 5x 15cm of MEFs, 2x 15cm ESCs, and 4x 15cm of reprogramming day 4 cells were used as starting material. Each ChIP was performed on a separate biological replicate expansion and duplicates were immunoprecipitated on separate days (Table 1). Cells were trypsinized and fixed with 1% formaldehyde in suspension for 10 min, rotating. Cross-linking was quenched with 0.14 M glycine for 5 min, cells were centrifuged at 300xg for 3 min, and pellets washed with cold 3x PBS. 25 million cells were resuspended in 1 ml lysis buffer (1% SDS, 50 mM Tris-HCl pH 8, 20 mM EDTA, 1x cOmplete (Roche, 4693132001) protease inhibitor), and sonicated on a Covaris S220 Focused-ultrasonicator with the following parameters: 21 cycles of 45 seconds ON (peak 170, duty factor 5, cycles/burst 200), 45 seconds OFF (rest) in 6-8°C degassed water. 25 ul aliquots were taken at 0, 10, and 21 cycles to check DNA fragmentation. Aliquots were diluted with 75 ul of H2O and incubated with 10 μg of RNase for 30 min at 37C. 1 ul of proteinase K (20 mg/ul) was added and sonication samples were incubated at 60 C overnight to reverse crosslinks. DNA was purified with phenol-chloroform extraction with phase lock tubes, precipitated with isopropanol, and run on a 1.5% agarose gel to ensure generation of 200-400 bp fragments.

**Table 1.**
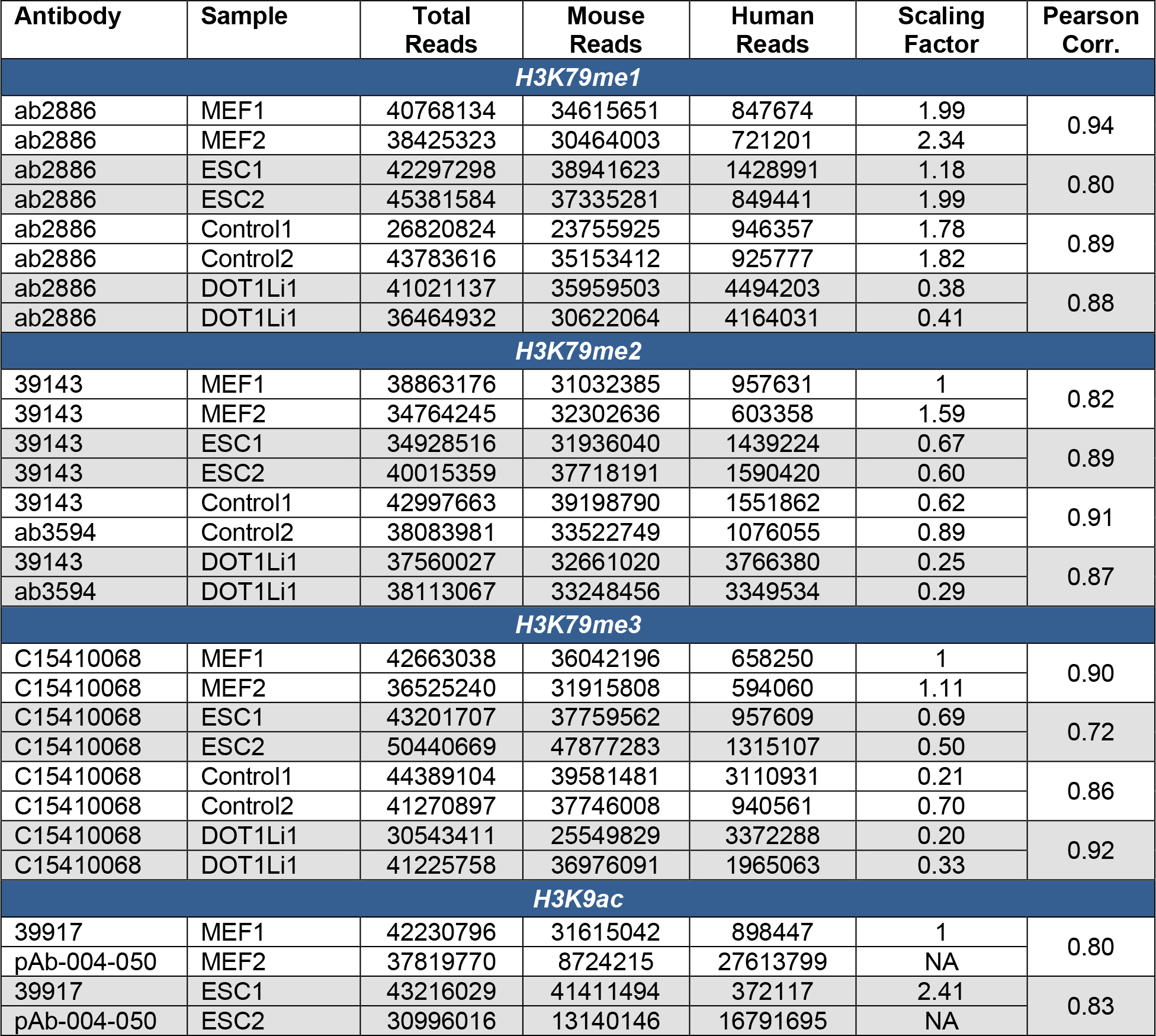
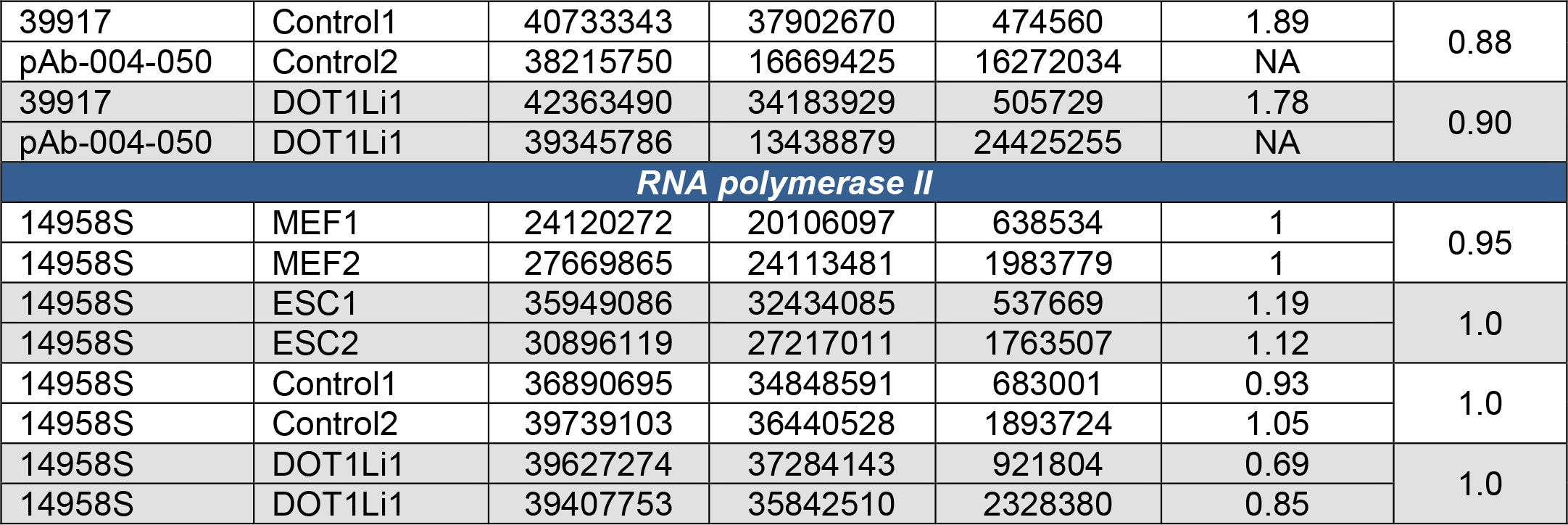
ChIP-Seq sample information.

| Antibody | Sample | Total Reads | Mouse Reads | Human Reads | Scaling Factor | Pearson Corr. |
| --- | --- | --- | --- | --- | --- | --- |
| H3K79me1 |  |  |  |  |  |  |
| ab2886 | MEF1 | 40768134 | 34615651 | 847674 | 1.99 | 0.94 |
| ab2886 | MEF2 | 38425323 | 30464003 | 721201 | 2.34 |  |
| ab2886 | ESC1 | 42297298 | 38941623 | 1428991 | 1.18 | 0.80 |
| ab2886 | ESC2 | 45381584 | 37335281 | 849441 | 1.99 |  |
| ab2886 | Control1 | 26820824 | 23755925 | 946357 | 1.78 | 0.89 |
| ab2886 | Control2 | 43783616 | 35153412 | 925777 | 1.82 |  |
| ab2886 | DOT1Li1 | 41021137 | 35959503 | 4494203 | 0.38 | 0.88 |
| ab2886 | DOT1Li1 | 36464932 | 30622064 | 4164031 | 0.41 |  |
| H3K79me2 |  |  |  |  |  |  |
| 39143 | MEF1 | 38863176 | 31032385 | 957631 | 1 | 0.82 |
| 39143 | MEF2 | 34764245 | 32302636 | 603358 | 1.59 |  |
| 39143 | ESC1 | 34928516 | 31936040 | 1439224 | 0.67 | 0.89 |
| 39143 | ESC2 | 40015359 | 37718191 | 1590420 | 0.60 |  |
| 39143 | Control1 | 42997663 | 39198790 | 1551862 | 0.62 | 0.91 |
| ab3594 | Control2 | 38083981 | 33522749 | 1076055 | 0.89 |  |
| 39143 | DOT1Li1 | 37560027 | 32661020 | 3766380 | 0.25 | 0.87 |
| ab3594 | DOT1Li1 | 38113067 | 33248456 | 3349534 | 0.29 |  |
| H3K79me3 |  |  |  |  |  |  |
| C15410068 | MEF1 | 42663038 | 36042196 | 658250 | 1 | 0.90 |
| C15410068 | MEF2 | 36525240 | 31915808 | 594060 | 1.11 |  |
| C15410068 | ESC1 | 43201707 | 37759562 | 957609 | 0.69 | 0.72 |
| C15410068 | ESC2 | 50440669 | 47877283 | 1315107 | 0.50 |  |
| C15410068 | Control1 | 44389104 | 39581481 | 3110931 | 0.21 | 0.86 |
| C15410068 | Control2 | 41270897 | 37746008 | 940561 | 0.70 |  |
| C15410068 | DOT1Li1 | 30543411 | 25549829 | 3372288 | 0.20 | 0.92 |
| C15410068 | DOT1Li1 | 41225758 | 36976091 | 1965063 | 0.33 |  |
| H3K9ac |  |  |  |  |  |  |
| 39917 | MEF1 | 42230796 | 31615042 | 898447 | 1 | 0.80 |
| pAb-004-050 | MEF2 | 37819770 | 8724215 | 27613799 | NA |  |
| 39917 | ESC1 | 43216029 | 41411494 | 372117 | 2.41 | 0.83 |
| pAb-004-050 | ESC2 | 30996016 | 13140146 | 16791695 | NA |  |
| 39917 | Control1 | 40733343 | 37902670 | 474560 | 1.89 | 0.88 |
| pAb-004-050 | Control2 | 38215750 | 16669425 | 16272034 | NA |  |
| 39917 | DOT1Li1 | 42363490 | 34183929 | 505729 | 1.78 | 0.90 |
| pAb-004-050 | DOT1Li1 | 39345786 | 13438879 | 24425255 | NA |  |
| RNA polymerase II |  |  |  |  |  |  |
| 14958S | MEF1 | 24120272 | 20106097 | 638534 | 1 | 0.95 |
| 14958S | MEF2 | 27669865 | 24113481 | 1983779 | 1 |  |
| 14958S | ESC1 | 35949086 | 32434085 | 537669 | 1.19 | 1.0 |
| 14958S | ESC2 | 30896119 | 27217011 | 1763507 | 1.12 |  |
| 14958S | Control1 | 36890695 | 34848591 | 683001 | 0.93 | 1.0 |
| 14958S | Control2 | 39739103 | 36440528 | 1893724 | 1.05 |  |
| 14958S | DOT1Li1 | 39627274 | 37284143 | 921804 | 0.69 | 1.0 |
| 14958S | DOT1Li1 | 39407753 | 35842510 | 2328380 | 0.85 |  |

Sonicated chromatin was centrifuged at 21xg at 6 C for 10 min, and supernatant was collected and quantified with Qubit DNA HS Assay Kit (ThermoFisher Scientific, Q32854). Chromatin was aliquoted ensuring the SDS remained in solution so that the concentration was exactly 0.1% after diluting 1:10 in Dilution Buffer (16.7 mM Tris-HCl pH 8, 0.01% SDS, 1.1% Trition-X, 1.2 mM EDTA, and 167 mM NaCl). H3K79me1/2/3 ChIPs started with 7.1 ug chromatin combined with 1/53 (134 ng) human spike-in chromatin generated from 293T cells as above^27^. H3K9ac and RNAPII ChIPs were immunoprecipitated with 13 ug chromatin with 1/53 (247 ng) human spike-in generated from human dermal fibroblasts for H3K9ac or 293T cells for RNAPII. 5 ug (or 5 ul if concentration was unknown) of the following antibodies were used H3K79me1 (Abcam, 2886), H3K79me2 (Active Motif, 39143), H3K79me2 (Abcam, ab3594), H3K79me3 (Diagenode, C15410068), H3K9ac (Diagenode, pAb-004-050 Rep 1, 293T spike in control), H3K9ac (Active Motif, 39917, Rep 2 dermal fibroblast control), and RPB1 NTD D8L4Y (Cell Signaling Technology, 14958S).

ChIPs were incubated overnight at 4 C with rotation and then Dynabeads were added for 2 hours. Dynabeads were pre-prepared as follows: 25 μL of Protein A (ThermoFisher Scientific, 10002D) and 25 μL of Protein G (ThermoFisher Scientific, 10004D) were combined and washed once in PBS, 0.02% Tween-20, once in dilution buffer, and then resuspended in an equal volume of dilution buffer. Antibody-bead complexes were washed twice for 5 min rotating at 4 C, in 1 ml of each of the following buffers: Low salt (50 mM HEPES pH 7.9, 0.1% SDS, 1% Triton X-100, 0.1% Deoxycholate, 1 mM EDTA pH 8.0, 140 mM NaCl), High salt (50 mM HEPES pH 7.9, 0.1% SDS, 1% Triton X-100, 0.1% Deoxycholate, 1 mM EDTA pH 8.0, 500 mM NaCl), LiCl (20 mM Tris-HCl pH 8, 0.5% NP-40, 0.5% Deoxycholate, 1 mM EDTA pH 8.0, 250 mM LiCl), and TE (10 mM Tris-HCl pH 8, 1 mM EDTA pH 8) using a magnetic rack. Beads were incubated with 250 μL elution buffer (50 mM Tris-HCl pH 8, 1 mM EDTA pH 8, 1% SDS) plus 200 ul TE with 0.67% SDS for 10 minutes at 65°C, shaking. 10 μg RNase A was added and incubated at 37 C for 30 minutes. Crosslinks were reversed overnight with 40 ug of proteinase K at 60 C. DNA was purified with phenol-chloroform extraction with phase lock tubes followed by isopropanol precipitation. DNA was resuspended in ultrapure H2O.

The entire ChIP was concentrated to 5 ul with a speedVac and used for library preparation with Ovation Ultralow System V2 (NuGEN, 0344) according to the manufacturer’s instructions. All reactions were performed in 0.5x the volume, and adapters were diluted 1/5 in water. Libraries were amplified for 10 cycles and signal was checked by running 10% on 1.5% agarose gel before the final bead purification. After bead purification, the remaining library was run on a 1.5% agarose gel, and an additional size selection was performed by cutting from 200-400 bp. The library was purified with the MinElute Gel Extration Kit (Qiagen, 28604). Quality was checked with DNA HS Assay Kit (ThermoFisher Scientific, Q32854) and Bioanalyzer3.0. Libraries were sequenced 1×50 on an Illumina HiSeq4000 at the NUSeq Core (Northwestern Feinberg School of Medicine).

### ChIP-Seq analysis

Reads were aligned to the mm9 genome assembly using Bowtie2^66^. Sam files were converted into bam files with Samtools-1.2^67^ view. Bam files were sorted with Samtools-1.2 sort. Reads that did not align to the mouse genome were isolated from sorted bam files with Sam-1.2 view -f4. These unaligned bam files were converted into fastq files with Samtools-1.2 bam2fq. The fastq files were then aligned to the human hg19 assembly and processed to sorted bam files as above. The number of reads that unambiguously aligned to the human genome were quantified from the sorted bam files with Samtools-1.2 flagstat, and used to calculate ChIP-Seq scaling factors (see Table 1).

Peaks were called using MACS2^68^ callpeak. All ChIP-seqs were processed with both default and broad peak calling algorithms, with p-values of 1e-3 and 1e-4 which were assessed in Integrative Genomics Viewer (IGV)^69^. H3K79me1/2/3 ChIP-seqs were called: --broad -p 0.0001, and H3K9ac narrow peaks were called with: -p 0.001. Peak files were annotated to the mm9 genome with HOMER^70^ annotatePeaks.pl. The output summary file was used to generate the percent genomic annotation stacked bar graphs. To determine genic lists for H3K79me ChIPs, genes with peaks within the gene body (5’UTR, exon, intron, 3’UTR, and/or TTS) were isolated, and ChIP-Seq sample replicates were overlapped using Venny2.0 to identify high confidence consensus genes. To identify cell-type specific H3K79me genic lists, the high confidence lists were overlapped using Venny2.0 and annotation was checked with Deeptools^71^ k-means clustering of heatmaps. To generate gene-associated lists for H3K9ac, genes with peaks within the gene body and/or the promoter were isolated, and ChIP-Seq sample replicates were overlapped using Venny2.0 to identify high confidence consensus genes. The high confidence consensus cell specific gene lists were compared in a 4-way Venn using the R package ggVennDiagram(x, label_alpha = 0, label = “count”)+ scale_fill_gradient. Gene ontology (GO) was performed with HOMER findMotifs.pl using the biological_process output or DAVID (https://david.ncifcrf.gov/tools.jsp) Functional Annotation Tool, Gene Ontology

GOTERM_BP_4 and GOTERM_MF_4.

Deeptools was used to create heatmaps and metaplots. First, normalized bigwig files were generated using the command bamCoverage with the following parameters: --binSize 10 - -scaleFactor (see Table 1) --ignoreForNormalization chrX. Matrices of normalized signal were created with the Deeptools computeMatrix command. In the case of H3K79me plots, the parameters were as follows: scale-regions -b 3000 -a 3000 --regionBodyLength 5000. For H3K9ac/RNAPII plots, the parameters were: reference-point --referencePoint TSS -b 500 -a 1500, or as listed. Deeptools was used to compare normalized signal of ChIP-Seq replicates by Pearson correlation with multiBigwigSummary -bs 1000. IGV was used to display example ChIP-seq tracks of normalized bigwig files. Reads over genomic coordinates were quantitated with BEDtools2.0^72^ multibamCov. Reads were normalized with the human spike-in scaling factor (see Table 1). In the case of H3K79me ChIP-seqs, scaled reads were further normalized to kb of gene length. To capture the DOT1Li-specific patterns in H3K9ac, we quantitated signal from TSS to +1 kb downstream (Fig. 6a). Traveling ratio was calculated by quantitating reads within the pausing region (from -30 to +300) and the elongation region (from +300 to transcription termination site) as previously described^36^. Reads were normalized to the length (kb) of the assessed region. The traveling ratio was calculated as the length normalized pause signal divided by the elongation signal, and averaged between the two quantitated replicate ChIP-Seqs per gene. All box and whiskers plots were generated in Graphpad Prism as: the box limits - 25^th^ to 75^th^ percentiles, the center line - median, and the whiskers - minimum and maximum values.

### Targeted DOT1L ChIP-qPCR

Lentiviruses were packaged that contained two small guide RNAs per target, as well as mCherry.

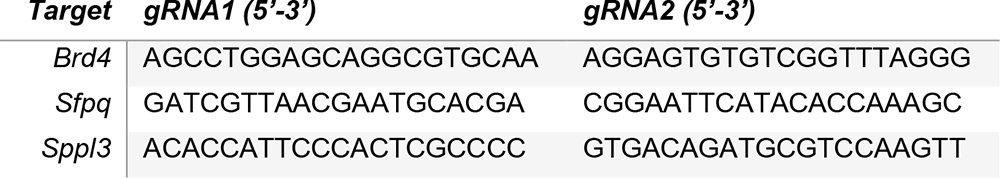

gRNA lentiviruses were titered with flow cytometry, and used to infect feeder-depleted the dCas9-DOT1L inducible ESC line at efficiency of at least 30%. Feeder MEFs, with and without 2 ug/ml dox induction, were added 6 hours post infection. Cells were cross-linked 48 hours post dCas9-DOT1L induction and processed for ChIP as in the ChIP-Seq Methods. ChIP for H3K79me2 started with 2 ug of chromatin, 1:13 spike-in 293T chromatin (153.8 ng), and 1 ug antibody (Active Motif, 39143). ChIP for H3K9ac started with either 7 ug (Rep 1) or 13 ug (Rep 2) of chromatin, 1:10 spike-in human dermal fibroblast chromatin (Rep1-700 ng or Rep2-1300 ng), and H3K9ac antibody (Rep 1-1.4 ug or Rep 2-5 ug) (Active Motif, 39917). Enrichment was calculated relative to the geometric mean of two genes highly enriched for both H3K79me2 and H3K9ac in human (h) cells (*RPLP1* and *EEF1A1*) from the spike-in chromatin. Primers are labeled by use: H3K79me2 (K79) ChIP, H3K9ac (K9) ChIP, Spike-in (SI) normalization, or non-targeted (NT) regions.

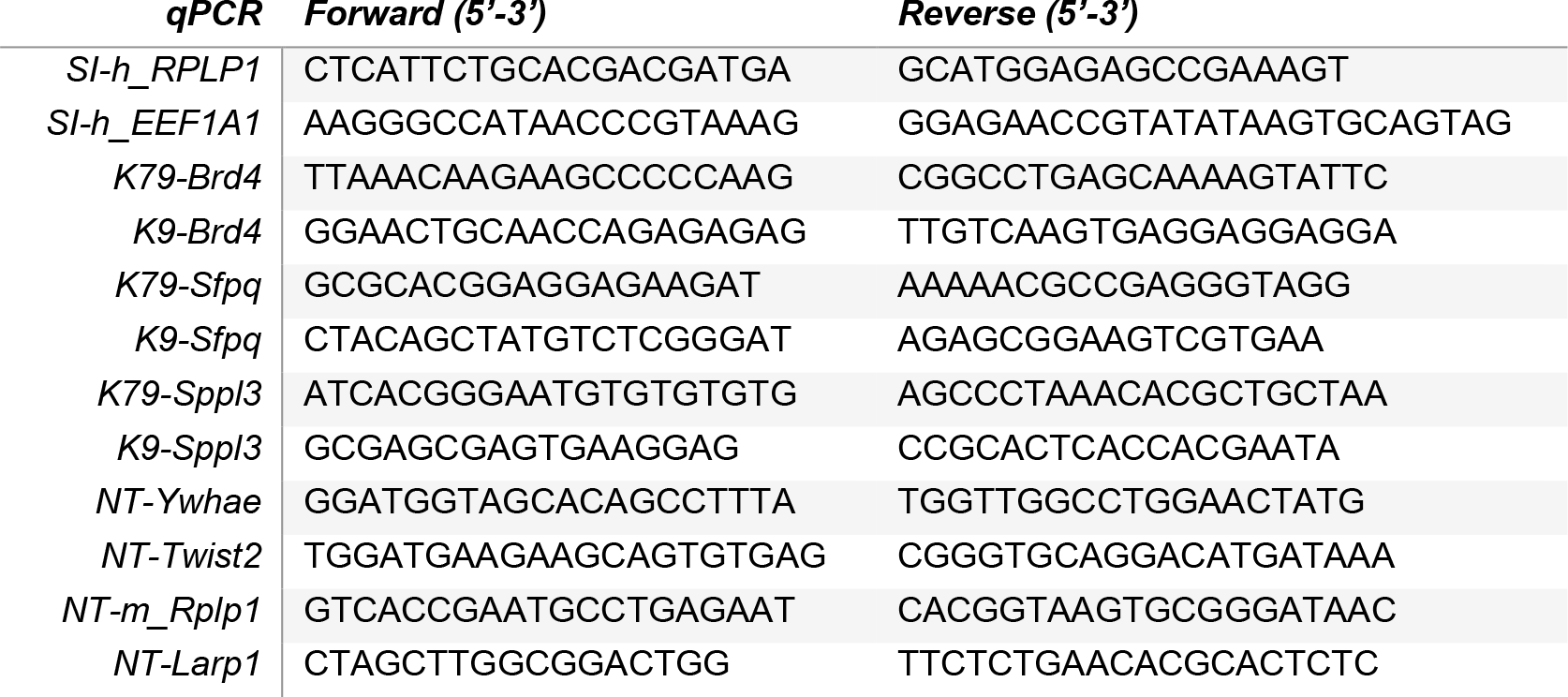

### Nascent RNA profiling

Cells were treated for 2 hrs and 50 min in 100 uM 5,6-Dichloro-1-β-D-ribofuranosylbenzimidazole (DRB) alone, and then media was changed to contain DRB with 5-ethynyluridine (EU) (Click Chemistry Tools, 1261 and Thermo Fisher Scientific, E10345). EU quantity was scaled based on cell number with a minimum concentration of 0.5 mM. After 10 min (3 hours total of DRB), the 0 timepoint was collected. For the transcriptional release timepoint, cells were washed twice in an excess of warm PBS to remove DRB, media with the scaled quantity of EU (and dox/drugs if in use) was replaced, and cells were collected 30 min after DRB removal.

Nascent RNA was assessed in MEFs, ESCs, and on day 6 of reprogramming with DMSO or SGC0946/DOT1Li using flow cytometry as previously described^73^. The click reaction was performed for 1 hour in click staining solution (100 mM Tris pH 8.5, 1 mM CuSO_4_, 2 mM AZDye-488 Azide (Click Chemistry Tools, 1275-1), and 100 mM ascorbic acid). An Attune Flow Cytometer (Flow Cytometry Laboratory, University of Wisconsin Carbone Cancer Center Support Grant P30 CA014520) or a BD Accuri C6 Flow Cytometer was used to analyze 10,000 cells per sample which were assessed using FlowJo software. Single cells were gated for analysis and per cell fluorescence was examined by exporting the scale values in FlowJo.

To profile nascent transcripts by qPCR, MEFs, ESCs, and 2D4 iPSCs were counted and harvested in TRIzol. Drosophila S2 cells that had been treated for 2 hours with 2 mM 5-EU were spiked in at a ratio of 1:20 (fly:mouse cells) for per-cell normalization. RNA was isolated with the RNA Clean & Concentrator-25 kit (Zymo Research, R1018). The Click-iT Nascent RNA Capture kit (ThermoFisher Scientific, C10365) was used according to the manufacturer’s instructions to pull down nascent EU-containing transcripts from 10 ug of starting total RNA. RNA was converted to cDNA from beads using qScript (Quanta, 95047) in 20 ul reactions. cDNA was diluted 1:4, and 1 ul was assessed by qPCR in 10 ul reactions. The Delta-Delta Ct was calculated relative to drosophila spike in, setting unreleased MEFs (treated for 3 hours with DRB) to 1. The following primers designed within introns at the indicated genic position downstream of the TSS were used:

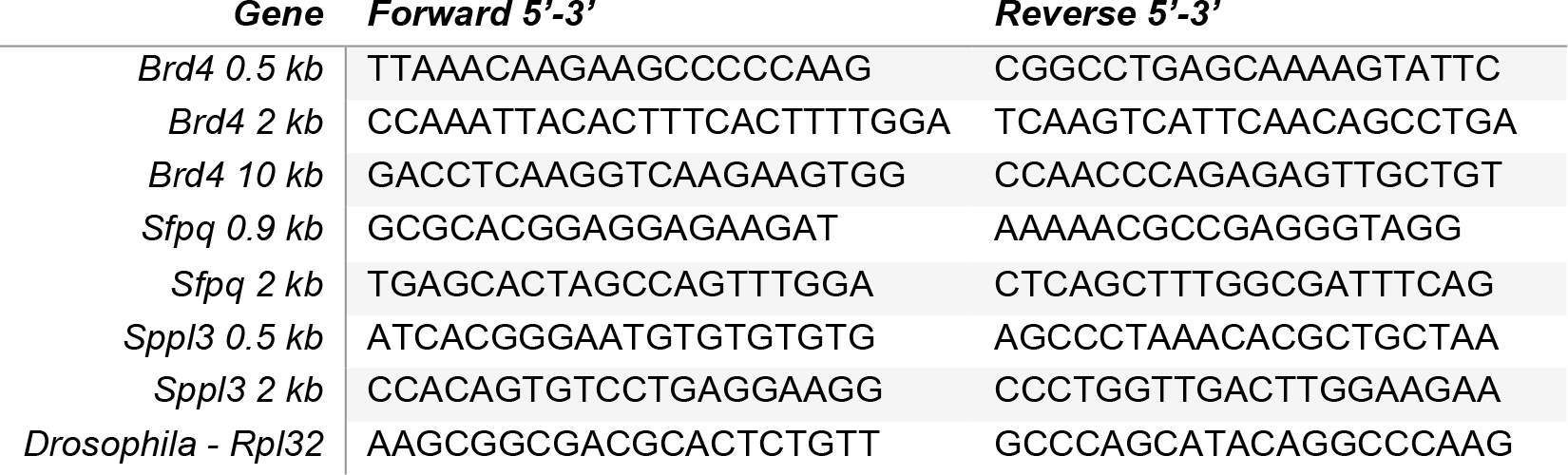

### Other Datasets

RNA-seq data GSE160580 was processed as previously described^25^. Other samples include:

**Other Datasets:** RNA-seq data GSE160580 was processed as previously described<sup>25</sup>. Other samples include:
| **Sample** | **GEO Sample** | **Cell type** | **Ref** |
| --- | --- | --- | --- |
| *D0_4E_K1* | SRX3372763 | Keratinocyte | SRP119979<sup>74</sup> |
| *D0_4E_K2* | SRX3372768 | Keratinocyte | SRP119979<sup>74</sup> |
| *WT_Naïve B cells (CD43-) Replicate 1* | GSM4121505 | Spleen B cells | GSE138909<sup>56</sup> |
| *WT_Naïve B cells (CD43-) Replicate 2* | GSM4121506 | Spleen B cells | GSE138909<sup>56</sup> |
| *WT_Naïve B cells (CD43-) Replicate 3* | GSM4121507 | Spleen B cells | GSE138909<sup>56</sup> |
| *Ctrl_E14.5_Dorsal Telencephalon_Rpl1* | GSM2526629 | Dorsal telencephalon | GSE95832<sup>75</sup> |
| *Ctrl_E14.5_Dorsal Telencephalon_Rpl2* | GSM2526630 | Dorsal telencephalon | GSE95832<sup>75</sup> |

## DATA AVAILABILITY

All ChIP-seq datasets have been submitted to the National Center for Biotechnology Information Gene Expression Omnibus database GSE190391.

## ACKNOWLEDGEMENTS

This work was supported by a UW-Madison Fall competition and Shaw Scientist award to the R.S. lab. We thank Dr. Roice Wille for aid in statistical analysis and script design, and members of the Sridharan lab for critical reading of the manuscript.

## AUTHOR INFORMATION

### Contributions

C.K.W. and X.Z. performed experiments, C.K.W completed bioinformatic analysis, S.A.H. performed mass spectrometry on an instrument located in the J.M.D. lab, C.K.W. and R.S. wrote the manuscript, R.S. acquired funding, and R.S. conceived and directed the project.

### Corresponding author

Correspondence to Rupa Sridharan.

