## Extended Data Figures for "Antagonistic H3K79me-H3K9ac crosstalk determines elongation at housekeeping genes to promote pluripotency"

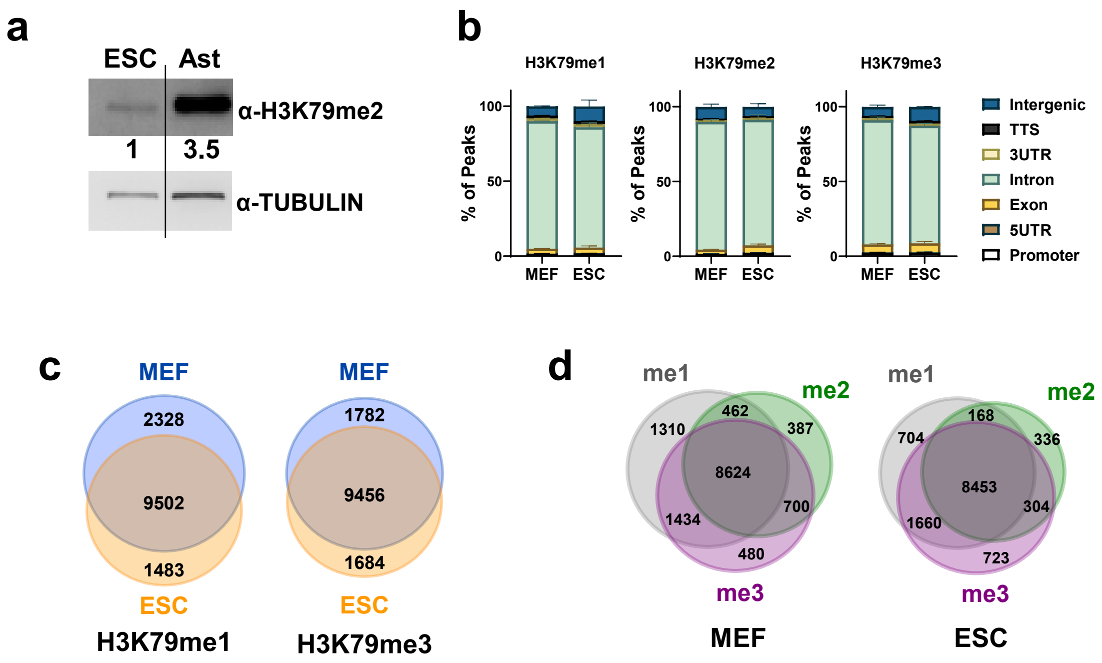


**Extended Data Figure 1.**

a. H3K79me2 levels in ESCs and astrocytes (Ast). Signal normalized to TUBULIN loading control and calculated relative to ESCs (numbers below). ESC panel also used presented in Wille et al., 2022.

b. Average percent of H3K79me1, me2, and me3 peaks per genomic annotation. TTS = transcription termination site and UTR = untranslated region. Data are mean + S.D. (n =2).

c. Overlap of genes with H3K79me1 (Left) or H3K79me3 (Right) peaks in MEFs and ESCs.

d. Overlap of genes with a significant H3K79me1/2/3 peak in MEFs or ESCs.

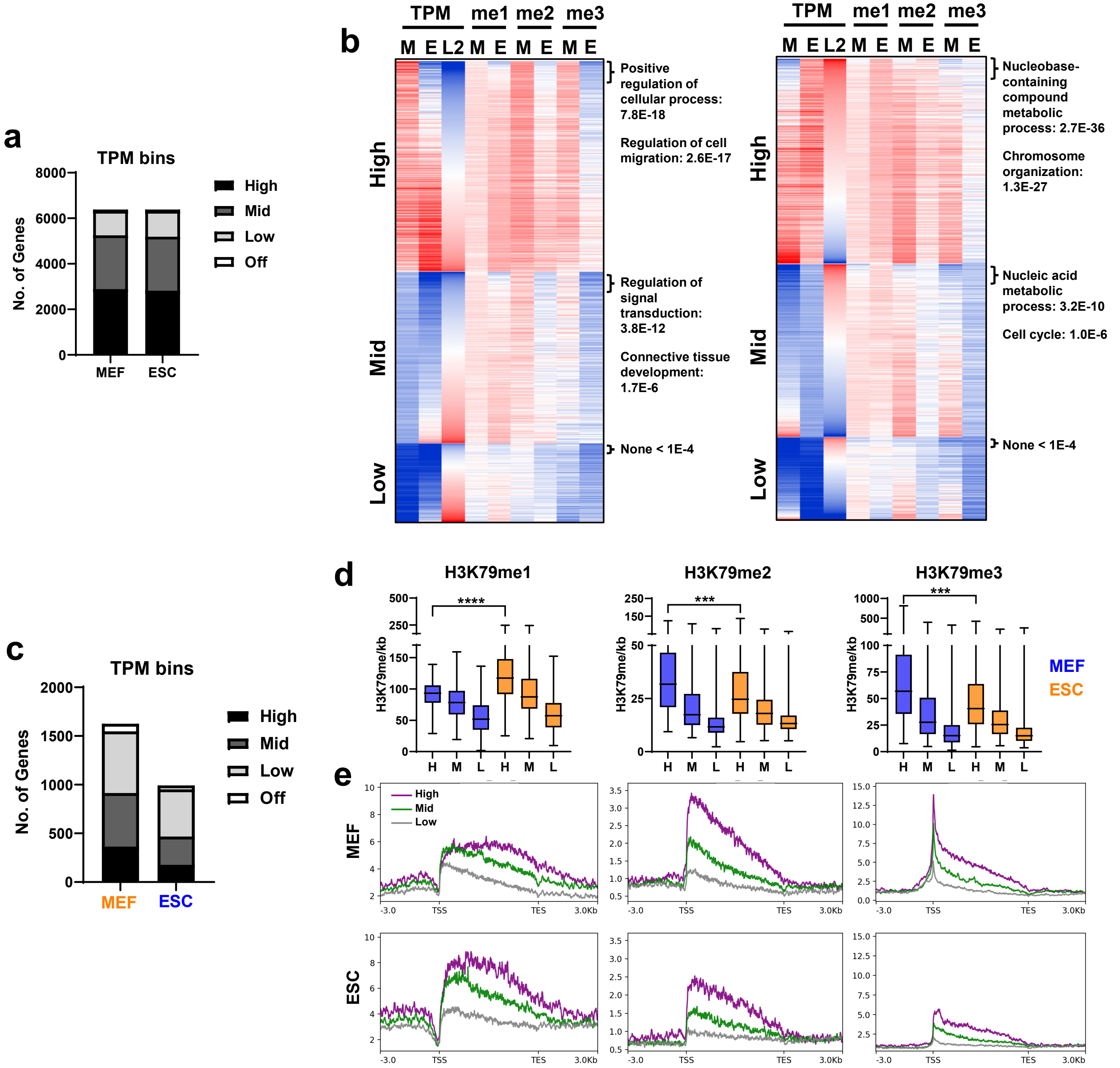


**Extended Data Figure 2.**

a. Number of genes per expression category with shared H3K79me2 peaks in ESCs or MEFs.

b. Heatmap of H3K79me (me1/me2/me3) reads per kb genebody, of genes with shared H3K79me2 peaks organized by MEF (Left) or ESC (Right) expression bins. Genes are sorted by Log2 fold change in ESC vs. MEF TPM per expression bin. Gene ontology (GO) of the top 10% most differentially expressed per bin is displayed.

c. Number of genes per expression bin with MEF- or ESC-specific H3K79me2 peaks.

d. Normalized H3K79me reads per kb genebody at High (H), Mid (M), and Low (L) expressed genes with MEF unique (blue) or ESC unique (orange) H3K79me2 peaks. ****P<0.0001 or ***P<0.001 by unpaired two-tailed t-test.

e. Metaplot of H3K79me1/2/3 per expression bin of genes containing MEF- or ESC-specific H3K79me2 peaks.

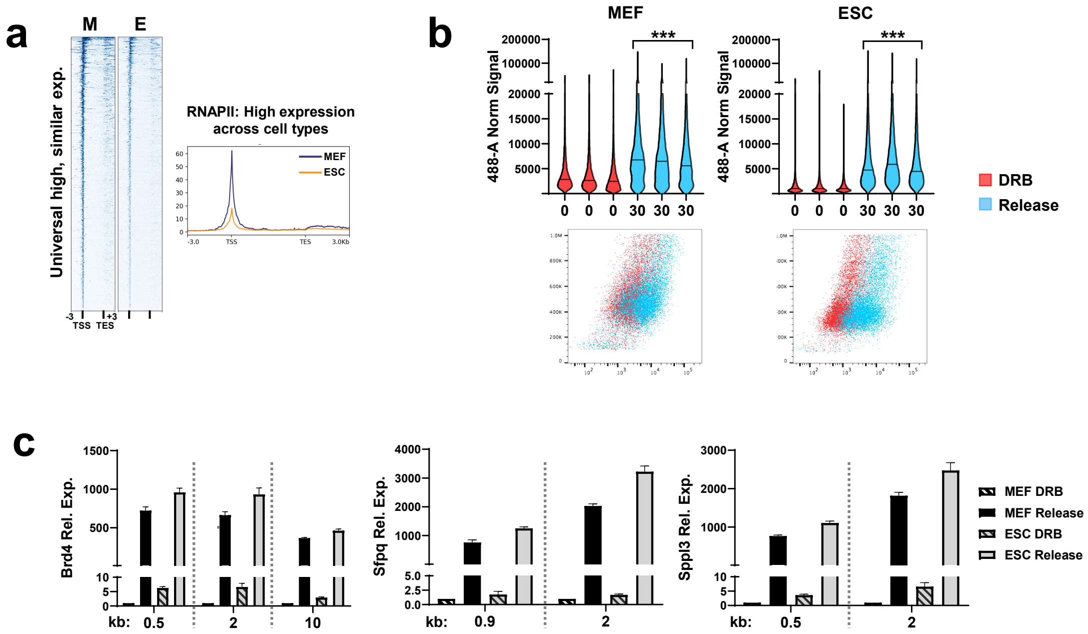


**Extended Data Figure 3.**

a. RNAPII heatmap (Left) and metaplot (Right) at H3K79me2 shared genes that have been filtered for universal high expression (same expression bin in MEFs, ESCs, keratinocytes, B cells, and brain in Fig. 2d), with similar total levels of expression (less than 2-fold difference in TPM) in ESCs (E) and MEFs (M).

b. Violin plot (Top) and representative scatterplot (Bottom) of EU-488 signal measured by flow cytometry. Red = DRB, Blue = Released 30 min. Individual replicates of each sample (n =3) are displayed (Top). ***P<0.001 by nested t test with a 0.95% confidence level. Note that MEFs have a greater background fluorescence in the presence of DRB.

c. Second biological replicate of relative nascent RNA RT-qPCR captured with 5-EU. Unreleased MEFs (DRB) set to 1. Data are mean + S.D. (n = 2 technical replicates).

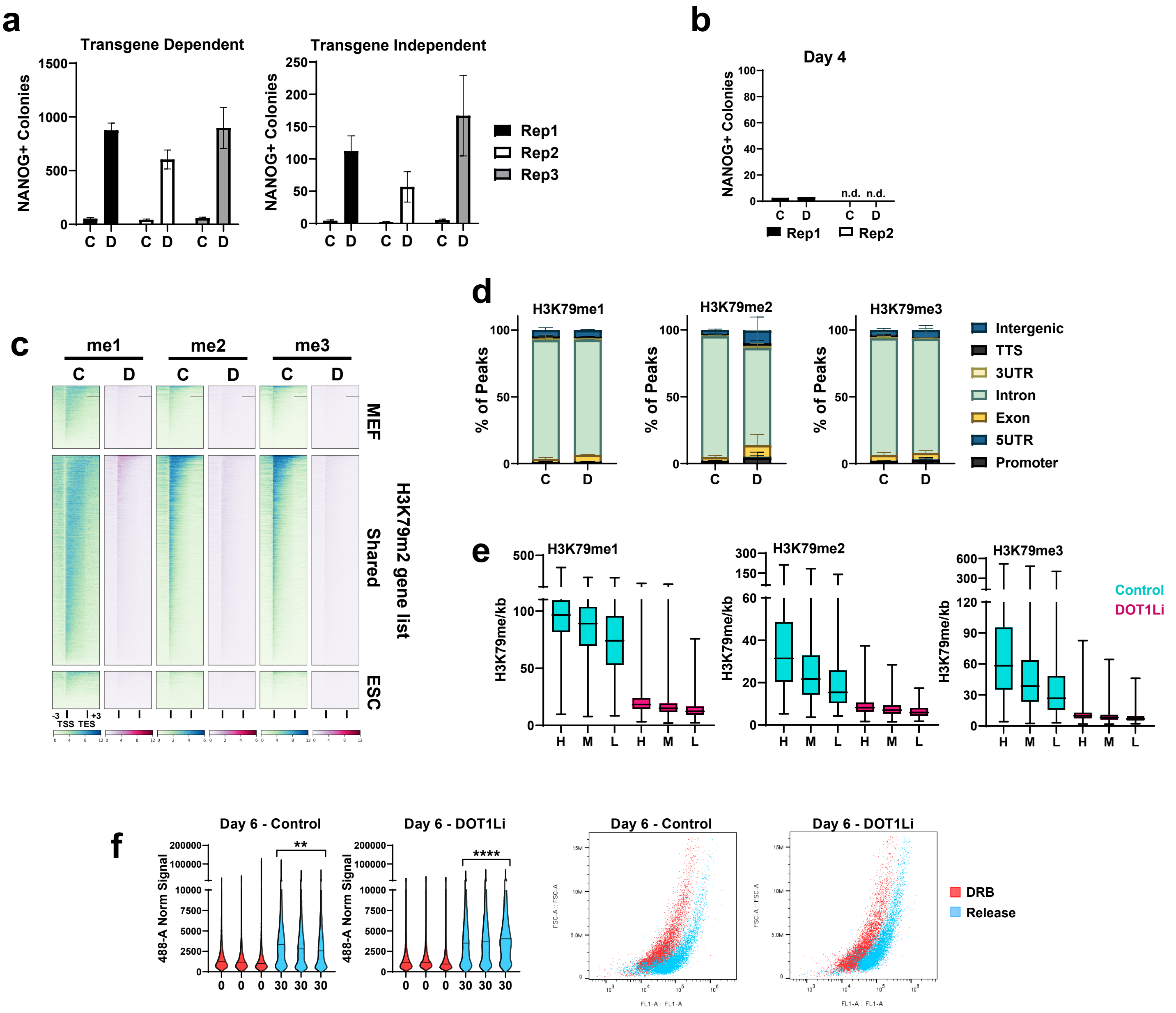


**Extended Data Figure 4.**

a. Transgene dependent (Left) and independent (Right) NANOG+ colonies of reprogramming cells treated with control (C) or DOT1Li (D) of each biological replicate. Data are the mean + S.D. (n = 3 technical replicates).

b. NANOG+ colonies on day 4 of reprogramming of the two independent biological replicates used for mass spectrometry analysis (Fig. 5). n.d. = not detectable. Data are mean + S.D. (n = 2 technical replicates).

c. Heatmap of H3K79me1, H3K79me2, and H3K79me3 signal on day 4 reprogramming in control (turquoise) and DOT1Li (pink) treated cells of MEF unique, ESC unique, shared genes with H3K79me2 peaks (Fig. 1a).

d. Average percent of H3K79me peaks per genomic annotation on day 4 of reprogramming in control (C) or DOT1Li (D) treated cells. TTS = transcription termination site and UTR = untranslated region. Data are mean + S.D. (n =2).

e. H3K79me1/2/3 normalized reads per kb genebody at shared H3K79me2 genes on day 4 reprogramming in control (turquoise) and DOT1Li (pink) treated cells. Genes separated into High (H), Mid (M), and Low (L) bins based on their expression in ESCs.

f. Violin plot (Left) and representative scatterplot (Right) of EU-488 signal measured by flow cytometry. Red = DRB, Blue = Released 30 min. Individual replicates of each sample (n =3) are displayed (Left). ****P<0.0001 or **P<0.01 by nested t test with a 0.95% confidence level.

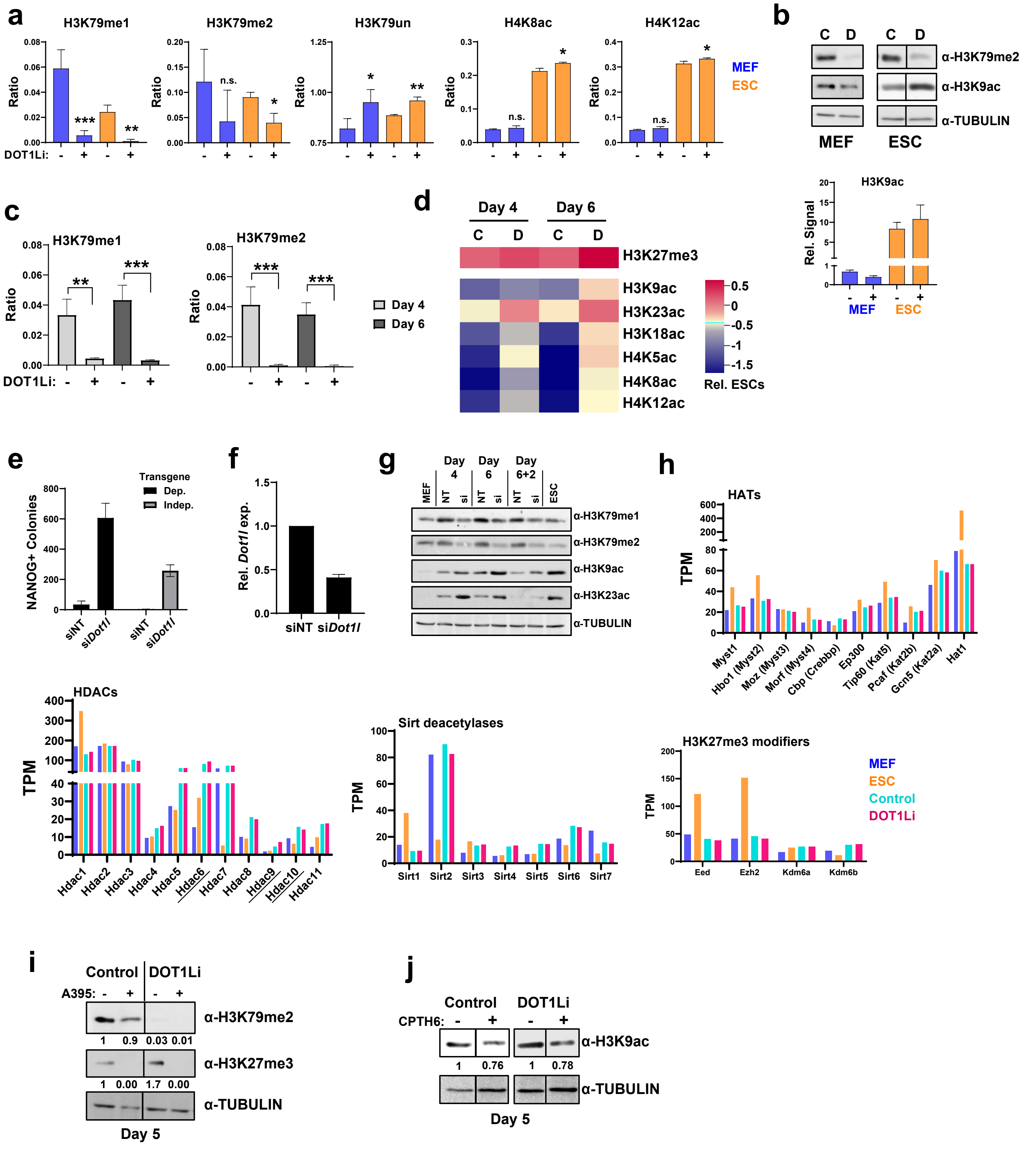
a
H3K79me1
H3K79me2
H3K79un
H4K8ac
H4K12ac
Ratio
DOT1Li:
MEF
ESC
b
C D C D
α-H3K79me2
α-H3K9ac
α-TUBULIN
MEF ESC
H3K9ac
Rel. Signal
c
H3K79me1
H3K79me2
Day 4
Day 6
d
Day 4 Day 6
H3K27me3
H3K9ac
H3K23ac
H3K18ac
H4K5ac
H4K8ac
H4K12ac
Rel. ESCs
e
Transgene
Dep.
Indep.
NANOG+ Colonies
siNT siDot1l siNT siDot1l
f
Rel. Dot1l exp.
siNT siDot1l
g
MEF NT si NT si NT si ESC
Day 4 Day 6 Day 6+2
α-H3K79me1
α-H3K79me2
α-H3K9ac
α-H3K23ac
α-TUBULIN
h
HATs
TPM
Myst1 Hbo1 (Myst2) Moz (Myst3) Morf (Myst4) Cbp (Crebbp) Ep300 Tip60 (Kat5) Pcaf (Kat2b) Gcn5 (Kat2a) Hat1
HDACs
Hdac1 Hdac2 Hdac3 Hdac4 Hdac5 Hdac6 Hdac7 Hdac8 Hdac9 Hdac10 Hdac11
Sirt deacetylases
Sirt1 Sirt2 Sirt3 Sirt4 Sirt5 Sirt6 Sirt7
H3K27me3 modifiers
Eed Ezh2 Kdm6a Kdm6b
MEF
ESC
Control
DOT1Li
i
Control DOT1Li
A395:
α-H3K79me2
1 0.9 0.03 0.01
α-H3K27me3
1 0.00 1.7 0.00
α-TUBULIN
Day 5
j
Control DOT1Li
CPTH6:
α-H3K9ac
1 0.76 1 0.78
α-TUBULIN
Day 5

**Extended Data Figure 5.**

a. Peptide ratio containing indicated modifications determined by mass spectrometry of MEFs (blue) or ESCs (orange) treated with and without DOT1Li for 4 days. Data are the mean + S.D. (n = 3-4). Significance determined relative to control treatment of the same cell type. ***P<0.001, **P<0.01, *P<0.05, or not significant (n.s.) P>0.05 by unpaired two-tailed t-test.

b. Top: Representative immunoblot of H3K9ac in MEFs and ESCs treated with Control (C) or DOT1Li (D) for 4 days. 10sec H3K9ac exposure shown for ESCs and 40 sec for MEFs. Bottom: Quantitation of relative immunoblot signal. Data are the mean + S.D. (n = 2).

c. Average peptide ratio containing the specified modification determined by mass spectrometry on day 4 (light) and 6 (dark) of reprogramming. Data are the mean + S.D. (n = 3-4). ***P<0.001 and **P<0.01 by unpaired two-tailed t-test.

d. Log2 fold change heatmap of select modifications in control (C) and DOT1Li (D) treated reprogramming cells relative to the ratio modified in ESCs in the second mass spectrometry biological replicate. Data are the mean (n = 3-4 technical replicates).

e. Transgene dependent (Dep. Black) and independent (Indep. Gray) NANOG+ colonies with control nontargeting (siNT) siRNA or si*Dot1l*. Extracts of this replicate assessed for histone modification levels in Extended Data Fig 5g. Data are mean + S.D. (n = 3 technical replicates).

f. Relative *Dot1l* transcript levels on day 4 of reprogramming with control nontargeting (siNT) or si*Dot1l*. siNT set to 1. Data are mean + S.D. (n = 2).

g. Immunoblot of histone modifications and TUBULIN control in MEFs, ESCs, and at the indicated reprogramming timepoints with control nontargeting (NT) and si*Dot1l* (si).

h. Histone modifying enzyme expression (TPM) in MEF (blue), ESC (orange), and reprogramming day 4 control (turquoise) and DOT1Li (pink) treatment. Samples altered more than 0.1 fold change in DOT1Li vs. control are underlined.

i. Immunoblot on day 5 of H3K79me2, H3K27me3, and loading control TUBULIN of control or DOT1Li treated cells, with A395 or vehicle beginning on day 0 of reprogramming. Relative quantitation shown below.

j. Immunoblot on day 5 of H3K9ac and TUBULIN of control or DOT1Li cells treated with CPTH6 or vehicle beginning on day 1 of reprogramming. Relative quantitation shown below.

**a**

| | Rep 1 | Day 4 | | Day 6 | | Rep 2 | Day 4 | | Day 6 | |
|---|---|---|---|---|---|---|---|---|---|---|
| | MEF | Control | DOT1Li | Control | DOT1Li | MEF | Control | DOT1Li | Control | DOT1Li |
| H3K9ac | -1.07 | -1.00 | -0.79 | -0.88 | -0.31 | -1.42 | -0.48 | -0.49 | -0.64 | -0.13 |
| H3K14ac | 0.11 | 0.08 | 0.20 | 0.04 | 0.23 | -0.22 | 0.28 | 0.28 | 0.08 | 0.19 |
| H3K18ac | -0.47 | -1.08 | -0.55 | -1.26 | -0.33 | -1.36 | -0.83 | -0.88 | -1.14 | -0.94 |
| H3K23ac | -0.32 | -0.43 | -0.06 | -0.36 | 0.06 | -1.02 | -0.17 | -0.03 | -0.41 | -0.10 |
| H3K27ac | n.d. | n.d. | n.d. | n.d. | n.d. | n.d. | -1.47 | -1.58 | -2.49 | -2.04 |
| H4K5ac | 0.07 | -1.35 | -0.42 | -1.62 | -0.30 | -1.94 | -0.55 | -0.63 | -1.25 | -1.10 |
| H4K8ac | -0.15 | -1.54 | -0.69 | -1.74 | -0.43 | -1.79 | -1.03 | -0.92 | -1.41 | -1.21 |
| H4K12ac | -0.28 | -1.29 | -0.56 | -1.51 | -0.40 | -2.08 | -0.97 | -0.88 | -1.57 | -1.27 |
| H4K16ac | -0.12 | -0.06 | 0.06 | -0.12 | 0.14 | 0.12 | 0.76 | 0.69 | 0.77 | 0.79 |

**b**

| | MEF | Control | DOT1Li | Control | DOT1Li | MEF | Control | DOT1Li | Control | DOT1Li |
|---|---|---|---|---|---|---|---|---|---|---|
| H3K4me1 | -0.44 | -0.03 | -0.05 | 0.10 | -0.04 | -0.30 | -0.47 | -0.14 | -0.26 | -0.11 |
| H3K4me2 | -0.18 | -1.14 | -0.40 | -1.10 | -0.16 | -1.22 | -1.05 | -0.83 | -1.00 | -0.99 |
| H3K4me3 | 0.50 | -0.49 | -0.02 | -0.49 | 0.23 | -0.61 | -0.54 | -0.54 | -0.43 | -0.77 |
| H3K9me1 | -1.10 | 0.19 | -0.43 | 0.25 | -0.53 | -0.33 | -0.14 | -0.16 | -0.15 | -0.08 |
| H3K9me2 | 0.10 | 0.05 | 0.09 | 0.04 | 0.10 | 0.07 | 0.15 | 0.11 | 0.14 | 0.03 |
| H3K9me3 | 0.22 | 0.05 | 0.10 | 0.06 | 0.06 | 0.62 | 0.13 | 0.22 | 0.18 | 0.17 |
| H3K18me1 | 1.23 | n.d. | -3.35 | -5.73 | -1.37 | 0.63 | -0.25 | -0.38 | -2.07 | -0.04 |
| H3K23me1 | 1.72 | n.d. | -2.86 | -6.20 | -0.88 | 1.43 | -2.72 | -0.27 | -3.85 | -1.99 |
| H3K27me1 | -0.98 | 0.54 | 0.14 | 0.46 | -0.65 | 0.05 | 0.04 | 0.13 | 0.04 | 0.02 |
| H3K27me2 | 0.03 | -0.25 | -0.14 | -0.20 | 0.06 | -0.01 | -0.11 | -0.17 | -0.04 | -0.05 |
| H3K27me3 | 0.79 | 0.08 | 0.19 | 0.09 | 0.62 | 0.20 | -0.61 | -0.35 | -0.26 | -0.03 |
| H3K36me1 | -0.70 | 0.21 | -0.06 | 0.32 | -0.61 | -0.05 | -0.42 | -0.32 | -0.33 | -0.30 |
| H3K36me2 | 0.30 | -0.67 | -0.38 | -0.87 | 0.16 | -0.30 | -0.20 | -0.22 | -0.30 | -0.41 |
| H3K36me3 | -0.40 | 0.97 | 1.02 | 0.85 | 0.28 | 0.65 | 0.47 | 0.56 | 0.56 | 0.59 |

**Extended Data Figure 6.**

Log2 fold change of detected H3.1 and H4 modifications relative to the ratio modified in ESCs. Modifications that DOT1Li makes cells more ESC like are boxed in blue. Modifications increased in MEFs relative to ESCs are boxed in red. Two biological replicates are displayed. Data derived from the mean (n = 3-4 technical replicates). Not detected = n.d.

a. Histone acetylation

b. Histone methylation

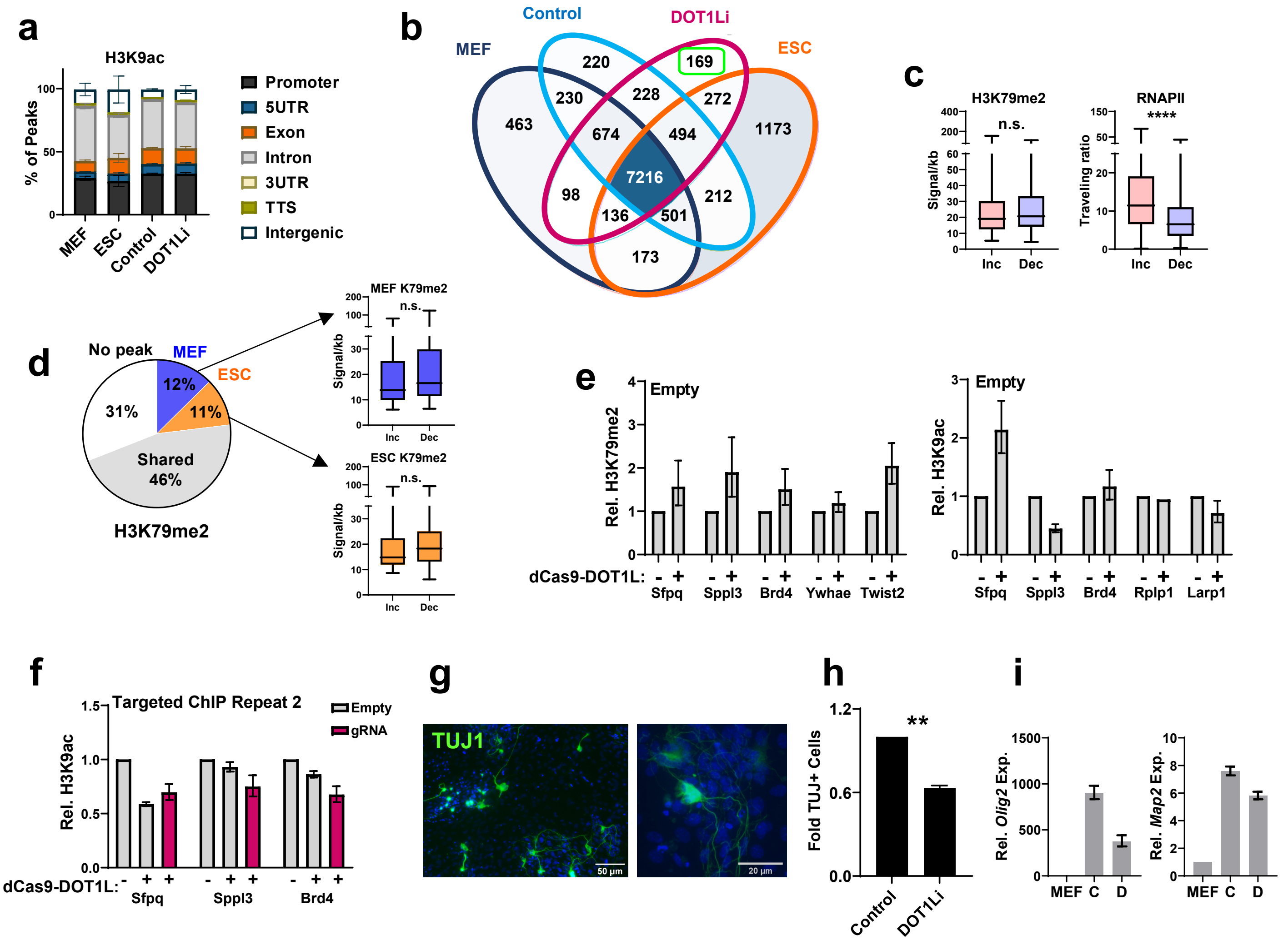


**Extended Data Figure 7.**

a. Average percent of H3K9ac peaks per genomic annotation. TTS = transcription termination site and UTR = untranslated region. Data are mean + S.D. (n =2).

b. Overlap of genes with genebody and/or promoter H3K9ac peaks. DOT1Li unique peaks boxed in green.

c. Genes with a shared H3K79me2 peak, in the top 10% with increased (Inc), compared to the bottom 10% with decreased (Dec), H3K9ac signal on day 4 of reprogramming in DOT1Li vs. Control (Fig. 6a) were isolated for analysis of: ESC H3K79me2 genebody signal per kb gene length (Right) and ESC RNAPII traveling ratio (TSS/genebody) (Left). Not significant (n.s.) P>0.05 by unpaired two-tailed t test

d. Genes with a MEF (blue)- or ESC (orange)-specific H3K79me2 peak, in top 10 and bottom 10% of relative H3K9ac signal on day 4 reprogramming in DOT1Li vs. Control (Fig 6a-b) were analyzed for H3K79me2 signal. Not significant (n.s.) P>0.05 by unpaired two-tailed t-test.

e. H3K79me2 (Left) and H3K9ac (Right) ChIP-qPCR, relative to spike-in chromatin, of cells infected with empty control lentivirus (no gRNA), with (+) and without (-) dCas9-DOT1L induction. The targets of the gRNA experiments (Fig. 7c) as well as non-targeted locations chosen for analysis.

f. Second biological H3K9ac ChIP-qPCR replicate of dCas9-DOT1L targeting, relative to spike-in chromatin. Cells were infected with empty control lentivirus containing no gRNA (gray), with (+) and without (-) dCas9-DOT1L induction, or infected with the indicated gRNA (pink) with (+) dCas9-DOT1L induction.

g. MEF transdifferentiation to induced neurons (iNeurons) with *Brn2*, *Ascl1*, and *Myt1l* (BAM) measured by TUJ1 expression.

h. TUJ1+ induced neuron cells on days 6-7 post BAM induction with control or DOT1Li treatment. Control treated cells set to 1. Data are the mean + S.D. (n = 2). **P<0.01 by unpaired t test.

i. Relative expression of neuronal factors in MEFs and on 7 of transdifferentiation treated with control (C) or DOT1Li (D). MEFs set to 1.
